# *In silico* discovery of compounds targeting NPSL2: a regulatory element in the human oncomiR-1 primary microRNA

**DOI:** 10.1101/2025.05.07.652683

**Authors:** Grace Arhin, Sarah C. Keane

**Affiliations:** Biophysics Program, University of Michigan, Ann Arbor, MI, USA 48109; Department of Chemistry, University of Michigan, Ann Arbor, MI, USA 48109

**Keywords:** virtual screening, microRNA, small molecule docking, nuclear magnetic resonance spectroscopy

## Abstract

NPSL2 is a stem loop element within the oncomiR-1 polycistronic primary microRNA (miRNA) cluster. NPSL2 is predicted to mediate conformational rearrangements within oncomiR-1 to regulate the biogenesis of certain miRNA elements within the cluster. The regulatory role of NPSL2 makes it a promising target for small molecule ligands, which we explored through structure-based small molecule ligand discovery. Starting with an NMR-derived ensemble of NPSL2 structures, we identified ligandable cavities within NPSL2 using RNACavityMiner. We then used molecular docking to screen the ZINC library against these cavities to identify potential small molecule hits. A sampling of commercially available compounds from the initial hits were purchased for experimental characterization by saturation transfer difference (STD) and heteronuclear single quantum coherence (HSQC) NMR spectroscopy. This approach led to the identification and characterization of at least eight compounds that bind preferentially within the internal loop of NPSL2. Importantly, this workflow, which combined virtual screening and experimental validation, can be used to identify small molecules that bind to any known RNA three-dimensional structure.

## INTRODUCTION

Mature microRNAs (miRNAs) are short non-coding RNAs (ncRNAs) that post-transcriptionally repress gene expression.^1,2^ MiRNA genes are transcribed as primary (pri) miRNAs which are subject to enzymatic processing by the Drosha/DGCR8 (DiGeorge syndrome critical region 8) complex, generating precursor (pre) miRNAs.^3,4^ These pre-miRNAs are subsequently processed by the Dicer/TRBP (transactivation response element RNA-binding protein) enzyme complex to produce mature miRNAs.^5^ The oncomiR-1 pri-miRNA is polycistronic, encoding six individual miRNA elements (miR-17, –18a, –19a, –20a, –19b, –92a).^6^ These miRNA elements are differentially processed by Drosha/DGCR8, leading to heterogeneous levels of the constituent pre-miRNA products.^6,7^ The oncomiR-1 cluster plays a prominent role in cancer development and progression including promoting cell proliferation, angiogenesis and metastasis with individual miRNAs regulating different cellular processes.^8,9^ Specifically, miR-92a has been reported to target the phosphatase and tensin homolog, a critical tumor suppressor gene, to promote metastasis and chemoresistance in non-small cell lung cancer,^10^ making it an attractive target for therapeutic intervention.

Secondary structure models of oncomiR-1, informed by chemical probing studies, reveal the presence of a small hairpin structure, NPSL2^6,11^ (**Fig. 1A**) within the 3ʹ-region of oncomiR-1 (between the pre-miR-19b and pre-miR-92a hairpins). NPSL2 hairpin formation disrupts the basal helix of pri-miR-92a,^6^ a required structural feature for Drosha/DGCR8 cleavage.^12,13^ Mutations to oncomiR-1 that eliminated the NPSL2 sequence reduced compaction in the 3ʹ-region of oncomiR-1 and exposed both miR-19b and miR-92a hairpins to RNase T1 digestion.^7^ Additionally, mutations to the NPSL2 sequence that disrupted NPSL2 tertiary contacts within the 3ʹ-region of oncomiR-1 enhanced pre-miR-92a expression.^7,14^ Interestingly, it is possible for NPSL2 to form an alternative conformation in which the hairpin structure is destabilized and the liberated sequence base pairs with nucleotides downstream of miR-92a (**Fig. S1**).^6^ This proposed conformational rearrangement would generate a basal helix for pri-miR-92a, making it a better substrate for Drosha/DGCR8 processing. As the NPSL2 structure is predicted to modulate accessibility of the Drosha/DGCR8 cleavage sites on pri-miR-92a, it is therefore an attractive drug target to control the levels of pre-miR-92a in cells. Small molecules that bind to and alter the NPSL2 hairpin stability may provide an indirect route to modulate the Drosha/DGCR8 mediated cleavage of miR-92a.

**Figure 1.**
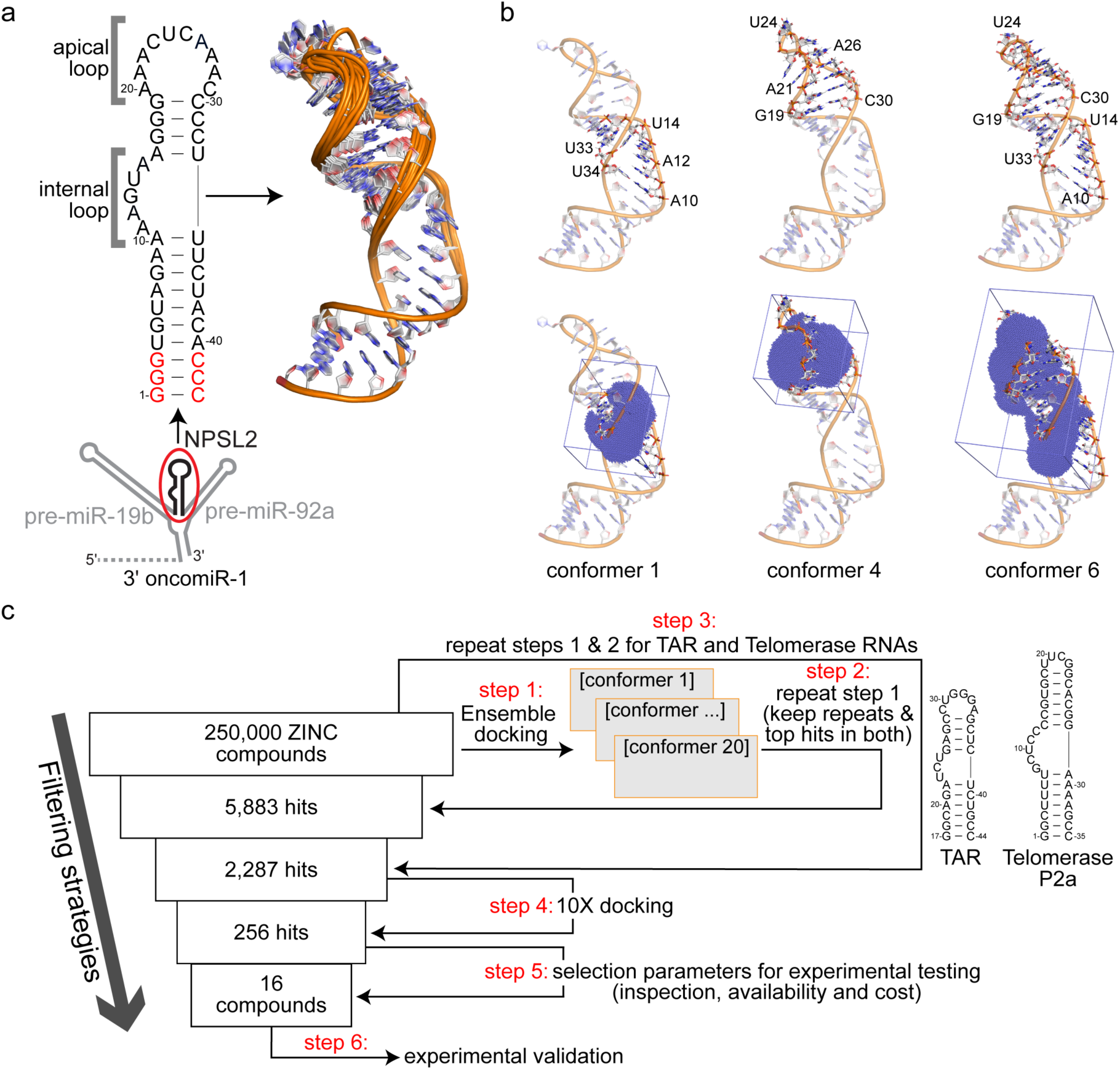
Structure based virtual screening approach. (a) Secondary structure of NPSL2 highlighting the 10-nt apical and 5-nt internal loops. The 3D NMR-derived ensemble is shown to the right. Beneath the NPSL2 secondary structure is a cartoon model of the oncomiR-1 3ʹ domain, highlighting the position of the NPSL2 hairpin. (b) 3D structures of conformers 1, 4 and 6 from the NPSL2 NMR ensemble showing their loop residues (top) and predicted binding volumes (in blue dots and a square box) determined from RNACavityMiner (bottom). (c) Schematic of the ensemble based virtual screening pipeline.

For functional and disease linked regulatory RNAs, RNA binding-small molecules ligands are often identified from a large pool of chemical libraries by high-throughput screens (HTS).^15^ However, structure based virtual screening (SBVS) can be used to screen a known RNA structure against even larger chemical libraries to rapidly identify compounds likely to bind the target RNA.^16^ A major downside to the SBVS approach is that biomolecular flexibility is generally disregarded. One strategy to address this limitation is to screen ligands against a structural ensemble rather than a single static structure.^17^ A second drawback of the SBVS approach is that the ligand search space is constrained to a predefined binding volume within the biomolecule. Therefore, in cases where a target binding site is unknown, blind docking is implemented, which is more expensive computationally and can reduce ligand search accuracy.^18^ To address this limitation, cavity searching algorithms can be used to identify potential ligand binding sites within the biomolecule, where docking can be focused.^19,20^

We previously determined the solution structure of NPSL2 using a combination of nuclear magnetic resonance (NMR) spectroscopy and small-angle X-ray scattering (SAXS).^21^ Our structural studies revealed that NPSL2 adopted a hairpin structure with a 10-nucleotide (nt) apical loop and a 5-nt internal loop (**Fig. 1A**). Because of the important role that NPSL2 structure is predicted to play in oncomiR-1 biogenesis, we sought to identify small molecule ligands that may interact with NPSL2 with a long-term goal of identifying molecules that function to inhibit production of pre-miR-92a.

In this study, the ligandability of the NPSL2 ensemble was assessed using RNACavityMiner.^19^ Through this analysis, we identified both the internal and apical loops of NPLS2 as sites that were likely to accommodate a ligand. We then developed an *in silico* protocol to virtually screen more than 250,000 compounds from the ZINC Library^22^ against the cavities identified within the NPSL2 ensemble, enabling the rapid identification of small molecule binders. We tested a subset of the *in silico* “hits” experimentally using both STD– and HSQC-NMR spectroscopy. This analysis led to the identification and characterization of several small molecule scaffolds that interact with NPSL2. Altogether, our findings highlight the importance of combining computational predictions with NMR-based approaches to readily identify and characterize RNA-ligand interactions with potential therapeutic applications.

## RESULTS

### Structure based virtual screening identifies compounds that bind NPSL2

We used RNACavityMiner^19^ to identify potential ligandable sites within the ensemble of NPSL2 structures.^21^ RNACavityMiner is a machine learning model that was trained on experimentally-determined RNA-ligand complex structures and predicts the likelihood that a given cavity in an RNA would accommodate a ligand.^19^ For the majority (60%) of the 20 NMR-derived conformers, RNACavityMiner predicted higher ligandability scores for the internal loop, whereas 30% of the conformers had a higher ligandability scores for the apical loop (**Fig. 1B** and **Fig. S2**). In roughly 10% of the conformers, RNACavityMiner predicted that both the apical and internal loops were likely to accommodate a small molecule ligand. This observed preference for ligand binding within the internal loop is consistent with the idea that it may be more favorable for ligands to bind smaller internal loop than the larger, more dynamic apical loop due to the entropic penalty paid for rigidifying a more flexible RNA.^23^

Using these identified binding volumes within NPSL2, we conducted a virtual screen of roughly 250,000 ZINC compounds^22^ against each NPSL2 conformer and identified 5,883 initial hits, i.e. compounds with the highest docking score (**Fig. 1C, steps 1 and 2**). To remove ligands that may be promiscuous RNA-binding molecules, we employed our virtual screening approach on two similarly structured RNAs. We first identified potential binding sites within the HIV-1 trans-activation response element (TAR) and telomerase P2a RNAs (7JU1^24^ and 2L3E^25^, respectively) (**Fig. S3** and **S4**). These RNAs were selected because they mimicked the size of either the internal or apical loop of NPSL2. We subsequently screened the entire ZINC library against the identified cavities for TAR and telomerase P2a RNAs. All highly scored compounds identified for TAR or telomerase P2a RNAs were eliminated from the hits identified for NPSL2 (**Fig. 1C, step 3**). Applying these selectivity conditions, we were able to reduce the initial hits identified for NPSL2 from 5,883 to 2,287 compounds.

Next, we docked these 2,287 ligand hits 10 times against the NPSL2 ensemble, generating 25 poses each time, keeping the top 10% of hits in each run (**Fig. 1C, step 4**). Collectively, these filtering strategies eliminated most compounds from the library and resulted in a pool of 256 hits. After inspection of the hits (**Fig. 1C, step 5**), we purchased 16 commercially available compounds for experimental characterization (**Fig. 1C, step 6**). Importantly, the SBVS hits selected for purchase contained about eight different scaffolds (**Fig. S5**) that might be useful for later structure activity relationship (SAR) studies. The computed docking free energies of the final 16 compounds are presented in **Table S1**.

### STD NMR screening of *in silico* hits against NPSL2

Our virtual screening workflow identified 256 hits, of which 16 were purchased for experimental testing. Many of the compounds had poor solubility in water and therefore needed to be prepared in a DMSO containing buffer. While DMSO helps to solubilize the small molecules, it is a chemical denaturant known to impact RNA structure at high concentration.^26^ Therefore, we first assessed the impact of DMSO on the NPSL2 structure by monitoring the imino proton region of the NPSL2 spectrum, which reports on base pairing within the RNA. We found that the secondary structure of NPSL2 was unaffected in buffer containing up to 5% (v/v) DMSO (**Fig. S6**).

With an upper limit of DMSO know, we next tested the solubility of each ligand in aqueous buffer containing 5% (v/v) DMSO. Of the 16 compounds we examined, eight compounds (Zn 588, Zn 699, Zn 273, Zn 735, Zn 126, Zn 043, Zn 591, Zn 513) were sufficiently soluble (at concentrations between 200-500 μM) to collect ^1^H-NMR spectra under these solution conditions (**Fig. S7-S14**). The other eight (Zn 918, Zn 981, Zn 668, Zn 711, Zn 091, Zn 079, Zn 347, Zn 504) were highly insoluble at concentrations below 200 μM in the buffer tested, with no observable signals in the ^1^H-NMR spectra (**Fig. S15-S22**). Ultimately, we excluded the eight poorly-soluble compounds from our ligand observed binding characterizations.

We acquired STD-NMR^27^ spectra (at a 1:20 RNA:ligand molar ratio) for the eight soluble compounds. As a ligand-observed experiment, STD can be used to rapidly identify ligands that bind to a biomacromolecule and characterize the binding epitope of the ligand within the complex (**Fig. 2a,b**).^27–29^ Our STD screening identified six compounds (Zn 588, Zn 699, Zn 591, Zn 043, Zn 735, Zn 126) that bound NPSL2 (**Fig. 2c**, **Fig. S23a-f**). Analysis of the difference spectra for Zn 513 and Zn 273 did not reveal interacting protons; indicating that these compounds may not be interacting with NPSL2 (**Fig. 2d**, **Fig. S23g,h**). We next examined whether our identified hits were selective towards NPSL2 by screening them against TAR and Telomerase P2a RNAs. The lead compounds appear to bind both TAR and Telomerase P2a RNAs, similar to NPSL2 (**Figs. S24**, **S25**). These observations make sense considering that small molecule binders of RNA often interact with generic secondary structural features such as loops, and bulges through H-bonding.^30^ Studies have shown that to increase selectivity in stem loop binders of RNAs, higher molecular weight compounds that increase the H-bonding interactions can be designed and tailored to an RNA target.^31,32^

**Figure 2.**
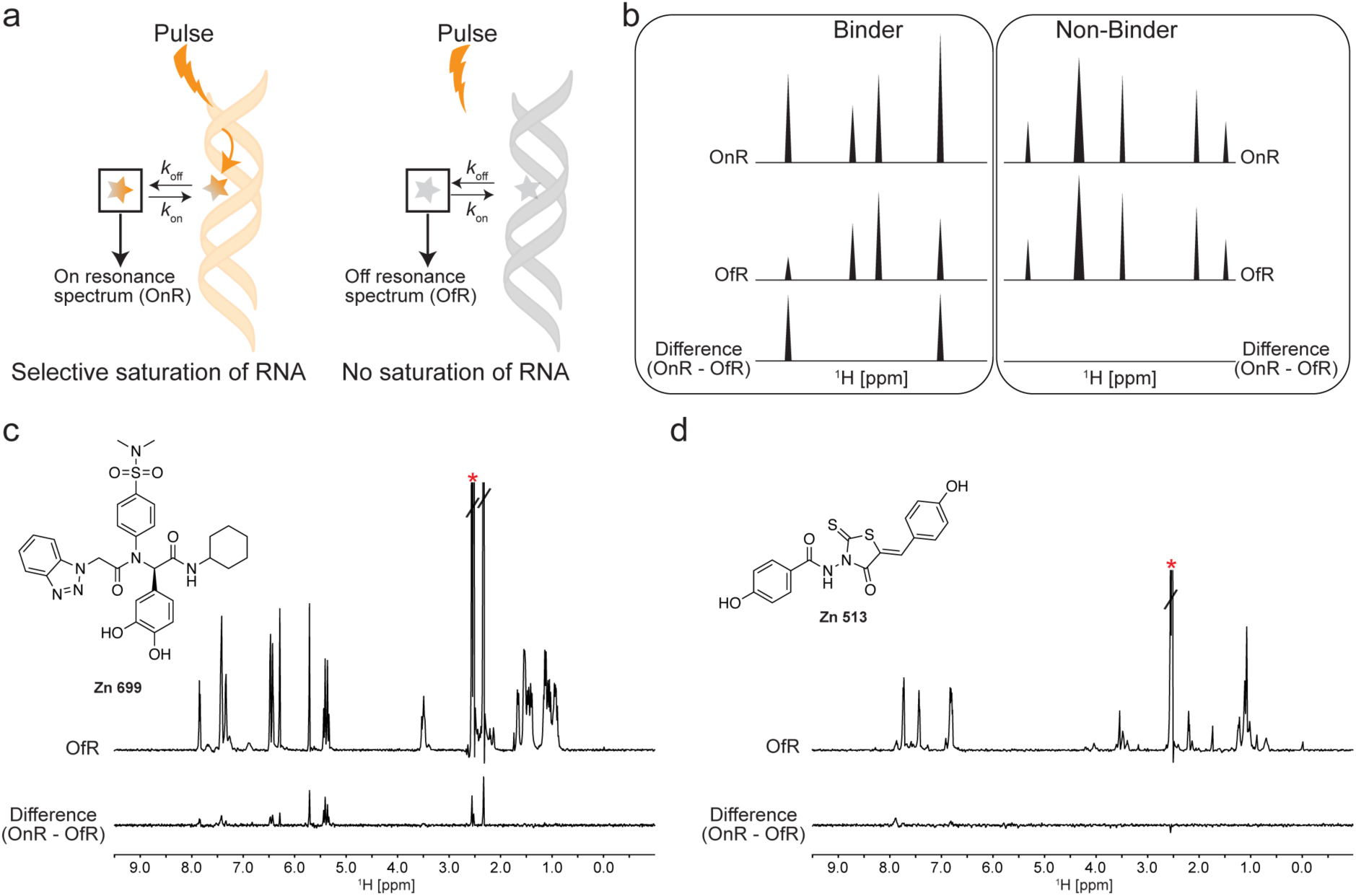
STD-NMR screening and hit identification. (a) Schematic of STD-NMR experiments. In the on-resonance experiment (OnR), the selective pulse saturates the RNA, indicated by the orange shading. If a ligand is bound, the cross-relaxation of the saturated RNA protons enhances the signals of nearby ligand protons. Additionally, an off-resonance spectrum (OfR) is recorded where there no saturation of the RNA. (b) Differences in the OnR and OfR spectra indicate binding. (c-d) Representative spectra of (c) a NPSL2 binder and (d) a NPSL2 non-binder. The top spectra are the ‘off” resonance spectra and the bottom spectra are the difference spectra. The red asterisk indicates the DMSO solvent peak.

### Evaluation of NPSL2 binding in closely related structures

Of the compounds analyzed by STD NMR spectroscopy, we noted three common scaffolds (**Fig. S26**). Within each scaffold, we sought to evaluate the impact of the different substituents on binding NPSL2. Compounds Zn 273, Zn 513, and Zn 735 all contain a thiazolidine like ring with a double bonded sulphur (S) and oxygen (O). For Zn 273 and Zn 513, the NH of the amide group is directly attached to the ring system while Zn 735 has a methylene group that connects the carbonyl group in the amide functionality to the thiazolidine ring. Our STD analysis revealed binding of Zn 735 to NPSL2 (**Fig. S23c**) while no binding by Zn 273 or Zn 513 was observed (**Fig. S23g,h**). These structural differences present different hydrogen bond donor/acceptor patterns which may contribute to the observed differences in binding with NPSL2 (**Fig. S26**). Zn 126 and Zn 043 share similar heterocyclic scaffolds (**Fig. S26**), and both interact with NPSL2 (**Fig. S23e,f**). However, due to the low signal:noise ratio of the Zn 126 difference spectrum, we were unable to analyze, and therefore compare, the binding epitopes.

Zn 588 and Zn 699 share a similar scaffold with differences in the R1 and R2 substituents (**Fig. 3a, Fig. S26**). We identified their binding epitopes to NPSL2 using STD NMR spectroscopy. We began by assigning the proton spectra of Zn 588 and Zn 699 in 100% DMSO (**Fig. S27a**,**b**; top spectra). Proton spectra of Zn 588 and Zn 699 in 95%/5% D2

**Figure 3.**
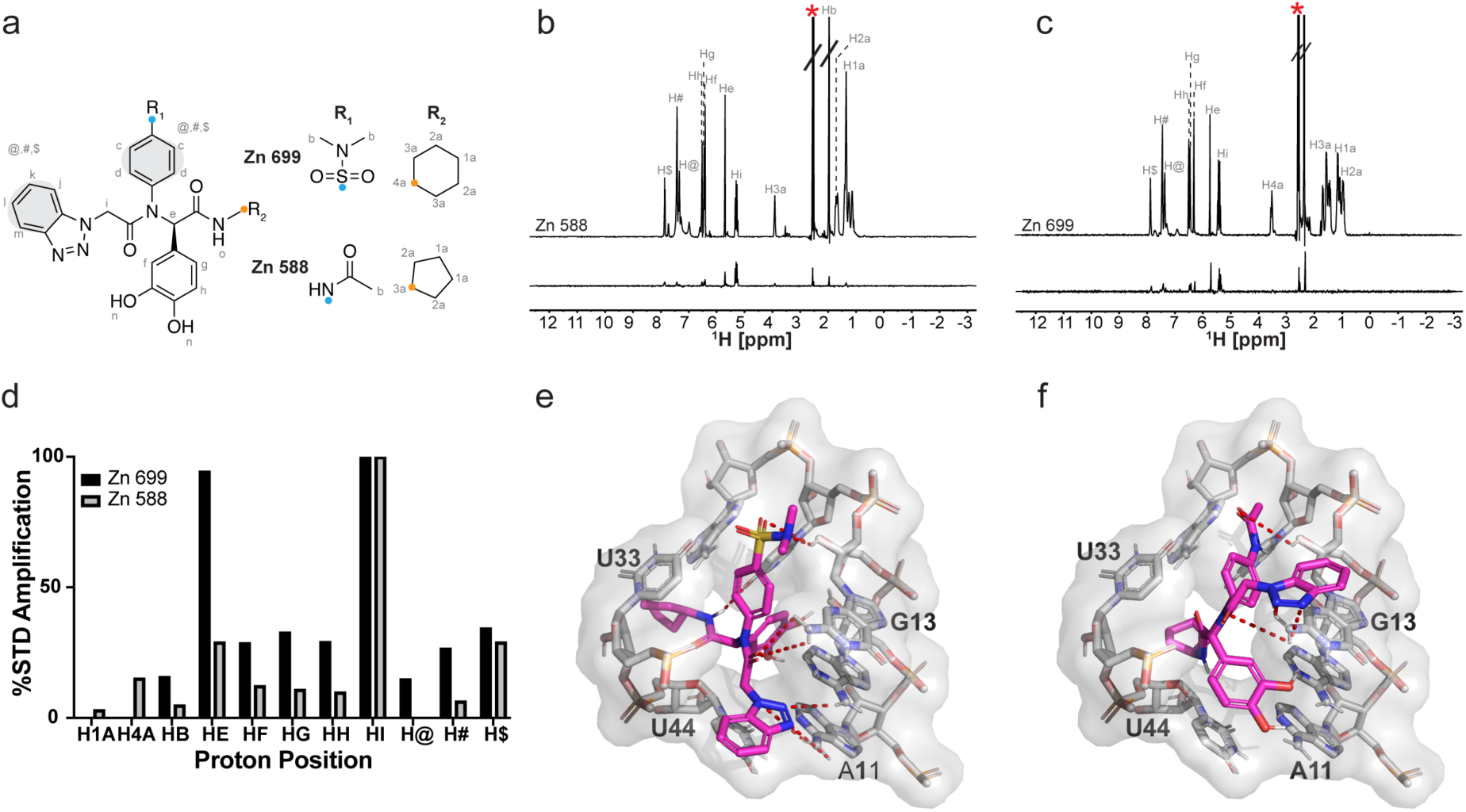
Binding characterization of hit compounds to NPSL2. (a) Shared scaffold between Zn 699 and Zn 588 showing the distinct R_1_ and R_2_ functional groups. (b-c) STD spectra of (b) Zn 588 and (c) 699 binding to NPSL2. (d) %STD amplification values of Zn 699 and Zn 588 of the distinct proton environments. (e-f) Molecular docking of (e) Zn 699 and (f) Zn 588 in NPSL2 showing their distinct poses.

O/DMSO exhibited line broadening of polar hydrogens peaks due to exchange with solvent. Furthermore, there was significant broadening of peaks corresponding to the protons on the benzene rings (between 7.2 – 8 ppm) preventing their precise assignment. Therefore, hydrogen-specific assignments for those regions could not be made and this group of hydrogens within the structures have been denoted as @, #, and $ (**Fig. S27a**,**b**; bottom spectra). We next analyzed the difference spectra of Zn 588 and Zn 699. Our results showed both compounds have distinct numbers of interacting protons (**Fig. 3b-d**). Notably, the proton peaks for R_2_ ring of Zn 588 (H1a and H3a) are weakly observed in the difference spectra indicating an interaction with NPLS2 (**Fig. 3b**). On the other hand, no protons are observed in the difference spectra for the R_2_ ring of Zn 699 indicating no contact with NPSL2 under the experimental conditions (**Fig. 3c**). We measured the percent STD amplification values for all interacting protons for Zn 588 and Zn 699 to enable comparison of proximity of the distinct protons to NPSL2 (**Fig. 3d**). We observed that for both compounds the proton (‘Hi’) in the main peptidomimetic chain is in closest proximity to NPSL2 when bound. All other peaks assigned amplification values relative to the ‘Hi’ proton show significance differences in peak intensities and proximities between the two compounds. Using these observed binding epitopes for both Zn 699 and Zn 588, we selected the most plausible binding pose from our docking simulations (**Fig. 3e**,**f**). Specifically, we evaluated which docked pose was most consistent with our observed STD interacting protons while also considering the computed score (binding energies) of the complex to ensure that the best pose was both consistent with STD results and energetically favored.

It is important to note that, while our STD analysis did not detect polar hydrogens and therefore could not be used as a direct measure of hydrogen-bonding interactions, we used the hydrogen-bonding interactions identified from the docked pose to explain the ligand orientation and which part of the ligand might be interacting with NPSL2. From this analysis, we found that pose 6 (the 6^th^ most energetically favored pose) for Zn 588 and pose 3 (the 3^rd^ most energetically favored pose) for Zn 699 best explained the observed STD data. These selected poses (**Fig. 3e**,**f**) provided a detailed visualization of ligand-NPSL2 interactions, revealing how the ligand interacts with specific NPSL2 residues while also explaining the observed STD effects. Interestingly, the identified poses were different for each ligand, reflecting the unique binding interactions each ligand has with NPSL2.

### HSQC-based screening of *in silico* identified hits with NPSL2

We next used chemical shift mapping experiments to explore interactions between our select SBVS hits (16 compounds) and NPSL2. This RNA-detected approach is complementary to the STD-based ligand-observed approach and offers additional advantages. Firstly, RNA-detected experiments provide a clear depiction of the regions within NPSL2 that are involved in ligand binding. Secondly, these experiments can be used to screen compounds that may not be amenable to ligand-observed experiments, particularly for ligands that have limited solubility and therefore weak ^1^H-NMR spectra.

2D ^1^H-^13^C HSQC spectra were recorded on an NPSL2 sample prepared with ^13^C/^15^ N-labeled adenosines both in the absence and presence of our compounds. We chose to isotopically label only adenosines to reduce spectral complexity while maintaining high sequence coverage. Adenosines account for 60% of residues in the internal and apical loops, which are the expected ligand interaction site(s). Our reference spectrum contained 5% DMSO to account for any subtle DMSO-induced perturbations to the NPSL2 spectrum. Notably, we observed minor perturbations across all adenosine resonances in NPSL2, including significant line broadening of the C8-H8 correlation of A21 (**Fig. S28**).

Upon addition of ligand to NPSL2, we monitored the ^1^H,^13^C HSQC spectrum for perturbations, which we recognized as either shifts in peak positions or changes in peak intensity, including the complete disappearance of peaks due to signal broadening. Specifically, for Zn 504, we observed a shift in peak position for A12 C8-H8 in addition to an increase in peak intensities for the C8-H8 peaks of A10, A11, A20, A21 and the C2-H2 of A10 and a slight decrease in peak intensity for C8-H8 of A12 (**Fig. 4a-c**). In the HSQC spectrum of NPSL2 in the presence of Zn 591, we observed quantifiable chemical shift changes for C8-H8 of A10, and A12. Furthermore, we observed a reduction in peak intensities for C2-H2 of A15 and C8-H8 of A12 and the complete disappearance of C2-H2 of A16 (**Fig. 4d-f**). Zn 591 was ^13^C-labeled and therefore exhibited resonances in the HSQC spectrum. We collected the HSQC spectra for Zn 591 alone, under the same experimental conditions, revealing minor perturbations in the ligand spectra when compared to the NPSL2 + Zn 591 spectra (**Fig. 4d,g**).

**Figure 4.**
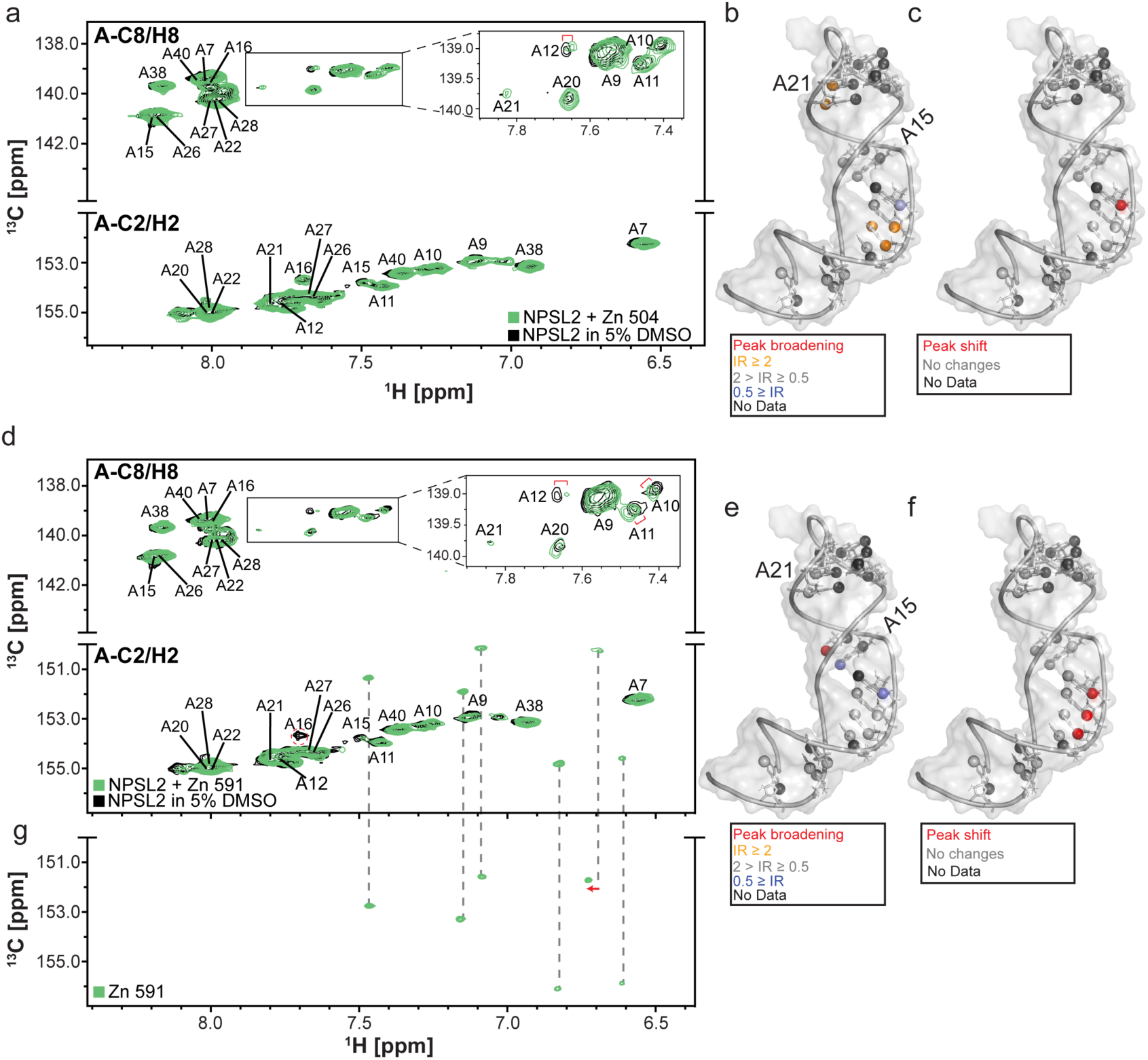
NMR chemical shift mapping experiments of Zn 504 (a-c) and Zn 591 (d-g) binding to NPSL2. Overlayed ^1^H-^13^C HSQC spectra of ^13^C/^15^N adenosine labeled NPLS2 (in 5% DMSO, black) with (a) Zn 504 and (d) Zn 591 (green). (b and e) Observed changes in intensity of NPSL2 resonances upon addition of Zn 504 (b) or Zn 591 (e) mapped onto the first conformer of the NPSL2 ensemble. (c and f) Perturbations arising changes in peak positions upon addition of Zn 504 (c) or Zn 591 (f) are colored red on the first conformer of the NPSL2 ensemble. (g) ^1^H-^13^C HSQC spectrum of Zn 591. The ligand signals are readily identifiable in the NPSL2 spectrum (d) with only minor perturbations (red arrow) relative to the free ligand state. Resonances exhibiting chemical shift perturbations are indicated with a red bracket. Resonances that exhibit complete signal broadening are marked with a red circle.

We observed similar perturbations for NPSL2 upon interaction with the following ligands; Zn 588, Zn 126, Zn 735 and Zn 668 (**Figs. S29-S32**), consistent with binding. Notably, these observed shifts were very small which may be the result of a small amount of ligand in solution interacting with NPSL2 due to poor solubility in our buffering system. For the remaining ligands (Zn 043, Zn 699, Zn 079, Zn 981, Zn 347, Zn 513, Zn 273, Zn 091, Zn 711, and Zn 918) we observed no significant perturbations to the NPSL2 spectrum (**Figs. S33-S42**). It should be noted that absence of significant perturbations in our NPSL2 spectrum upon addition of these ligands does not primarily signify the absence of a binding event. For example, Zn 043 and Zn 699 bound NPSL2 in our STD-NMR analysis thus the absence of chemical shift perturbation may be an indication of a weakly binding ligand.

### Experimental “hits” do not influence the thermal stability of NPSL2

To complement our insights obtained from NMR binding assays which indicated binding for Zn 591, Zn 504, Zn 588, Zn 688, Zn 699, Zn 735, Zn 126 and Zn 043, we evaluated the impact of these compounds on the thermal stability of NPSL2. Using CD thermal denaturation assays, we determined the melting temperature (*T*_*m*_), enthalpy (Δ*H*) and entropy (Δ*S*) associated with the NPSL2-ligand complexes (**Table S2, Fig. S43**). Despite evidence of ligand binding in other assays, we observed negligible changes in the melting temperature of NPSL2 upon addition of these ligands. These small changes may be a result of the ligands’ weak affinity for NPSL2. Furthermore, ligand binding may be affecting stability of NPSL2 at specific local regions (such as loops) rather than inducing any large-scale structural rearrangements to accommodate the ligands.

## DISCUSSION

Prior studies have posited that NPSL2 may regulate the Drosha/DGCR8 cleavage of miR-92a within oncomiR-1.^6,7^ To identify small molecule ligands that may bind NPSL2, we successfully implemented a structure-guided approach, integrating molecular modelling with experimental characterization. This integrated approach leverages the efficient and rapid power of computing with the accuracy of experiments.

There are a number of experimental screening approaches to identify RNA binding small molecules including microarray screens,^33^ fluorescent indicator assays,^34^ and NMR-based approaches,^35,36^ and each technique has strengths and limitations. These experimental techniques do not require tertiary structure information of the target RNA, making them suitable for screening small molecules against RNAs with unknown structures. However, this also means they provide little, if any, molecular level details of the RNA-small molecule interactions. These molecular insights are invaluable for rational small molecule design.^15^ These experimental approaches can be used independently to assess ligand binding but can also be used in combination with structure-based screening (virtual screening) techniques. By integrating SBVS with STD-NMR spectroscopy, ligands targeting the U-rich region of human telomerase P2b RNA have been identified.^28^ Additionally, SBVS has been used in combination with a fluorescence indicator assays and UV-melting studies to discover five small molecule ligands that bound MALAT1 triplex, with one compound demonstrating antiproliferative activity against multiple myeloma.^37^ Recent studies identified ligands targeting a large internal loop within the cis-regulatory epsilon RNA of the Hepatitis B virus using a combination of SBVS, molecular dynamics simulations, and fluorescent indicator assays, providing valuable insights into the RNA-small molecule interaction.^38^

Our screening approach integrates SBVS with experimental characterization using both STD NMR and chemical shift mapping experiments. This approach allowed us to obtain complementary information, monitoring changes in both the RNA and the ligands, to provide a more complete picture of the RNA-ligand interactions. Structural studies of small molecule ligands interacting with RNAs reveal a tendency for the ligands to form base stacking and Hydrogen bond interactions with residues in and around bulges or internal loops.^15,28,39–41^ Our docking studies revealed that ligands tended to form base stacking and hydrogen bonding interactions with residues within the internal loop of NPSL2. Consistent with the docking studies, our chemical shift mapping experiments revealed perturbations to NPSL2 resonances particularly in and around the internal loop and the apical loop.

While our structure-guided approach can be very useful in the identification of ligands targeting RNAs, there are several important limitations to note. One significant challenge is that several active compounds may be flagged as ‘non-hits’ during initial virtual screening stage which would result in their exclusion from experimental testing. Therefore, the overall success of our approach is dependent on the accuracy of the scoring function used in the virtual screening, which may not always capture the complexity of the biological system being studied. Additionally, in our virtual screening approach we attempted to identify compounds selective to NPSL2 by screening our library against other similarly structured RNAs and eliminating probable binders from our NPSL2 hits. Notably, in our experimental characterization we observed that the identified hits, which we expected to be selective for NPSL2, interacted with similarly structured RNAs. Thus, ligand selectivity may not best be adequately addressed at the virtual screening stage and thus our approach may have inadvertently eliminated some high affinity hits for NPSL2. Furthermore, while NMR-based approaches are useful for validating and characterizing *in silico* hits, a key limitation is the requirement for higher sample concentrations. These sample requirements make it extremely challenging, if not impossible, to study compounds that are poorly soluble in the aqueous buffer system.

We further note that, while we incorporated RNA structural diversity by screening against an NMR ensemble, that ensemble was relatively well-defined and did not sample a very large conformational space. It is possible that NPSL2 does in fact sample additional lowly-populated conformations not represented in the NMR ensemble. Therefore, using a more diverse ensemble that captures the different conformational states of the RNA, could significantly enhance hit identification, as was demonstrated for TAR.^42,43^

Despite these limitations, our approach has aided the identification and characterization of several compounds that bind to NPSL2, highlighting the powerful synergy between molecular modeling and NMR in the identification RNA-binding ligands. The initial hits from our screen serve as an excellent starting point for the rational design of compounds with increased affinity and selectivity towards NPSL2. Furthermore, additional experiments are required to investigate the impact of ligand binding to NPSL2 on the biogenesis of miR-92a in oncomiR-1 both *in vitro* and *in cell*. Our work provides a useful platform for future explorations and the methodological approach described herein will impact medicinal chemistry approaches especially for functionally relevant RNAs.

## METHODS

### Cavity identification and docking site preparation

We carried out cavity identification using RNACavityMiner.^19^ The coordinates of the NPSL2 NMR ensemble (7UGA, 20 structures)^21^ were split and converted into individual pdb files. Next, the main script (miner_grid.sh) of the RNACavityMiner package which takes as input the coordinates of the RNA was run on the individual NPSL2 conformers. The script outputted a cavity_pruned file (a .sd file) for each conformer which contains place holder atoms for the predicted binding site. Next, rbcavity from the rDock programs was run to generate the docking sites (.as) file which contains the cavity volume. PyMOL molecular visualization software as used to map these cavities onto their corresponding conformer to visualize binding site (**Fig. S2**). The same approach was used to identify cavities for TAR (7JU1)^24^ and telomerase P2a (2l3E)^25^ RNAs (**Fig. S3, Fig. S4**).

### Virtual screening

Using the rDock^44^ computational docking package, we screened 250,000 compounds from the ZINC library.^22^ The coordinates of all 20 conformers of the NMR ensemble of NPSL2 (7UGA)^21^ were individually docked (10 poses each) against the ligand library. Docking was focused on the druggable pockets (identified in the grid file from RNACavityMiner, above). A two-phase docking protocol was carried out. In the first phase, the entire library was docked against the NPSL2 ensemble (20 conformers) twice, allowing 10 possible poses per ligand. Hits were identified as compounds that were three standard deviations lower than the mean binding energy (score) in the ligand library. Our selection was based on top hits for each conformer as well as repetitive hits in both docking runs. Additionally, the library was docked against other RNA ensembles with similar structural motifs (telomerase P2a RNA [2l3E]^25^ and TAR RNA [7JU1]^24^), exactly as described above as an added selectivity filter to remove promiscuous binders. Hits identified for TAR and telomerase RNAs were eliminated from our NPSL2 hits.

In the second phase, the hits from phase one were each docked 10 times (10x docking) against the NPSL2 ensemble, allowing 25 possible docking poses per ligand. We then selected as hits, compounds that repeatedly docked the NPSL2 ensemble in each run as well as top 10% hits from every single run as our final hits for NPSL2 (256 compounds). Finally, we purchased 16 top commercially available compounds from eMolecules for experimental binding studies against NPSL2.

### Preparation of DNA duplex for transcription

The DNA templates for NPSL2, TAR, and telomerase P2a RNAs were purchased from Integrated DNA Technologies. Each template was 2ʹ-*O*-methoxy modified at the two 5ʹ most residues to reduce N + 1 product formation by T7 RNA polymerase.^45^ The partially duplexed DNA required for transcription was prepared by annealing each DNA template with a 17-nt DNA corresponding to the T7 promoter sequence (5ʹ-TAATACGACTCACTATA-3ʹ).

### RNA transcription

RNA was prepared by *in vitro* transcription with T7 RNA polymerase as previously described.^21^ Briefly, DNA template, ribonucleoside triphosphates (rNTPs), magnesium chloride (MgCl_2_), yeast inorganic pyrophosphatase (New England Biolabs), and dimethyl sulfoxide (DMSO)^46^ were mixed with in house prepared T7 RNA polymerase. The transcription reaction was incubated at 37 °C with shaking at 75 rpm. After 3 h, the reaction was quenched with a solution of 7 M urea and 500 mM ethylenediaminetetraacetic acid (EDTA), pH = 8.5. RNA was purified from the crude transcription reaction by denaturing polyacrylamide gel electrophoresis. The quenched transcription reaction was loaded onto a 16% preparative scale denaturing polyacrylamide gel with the addition of 16% glycerol (50% v/v) and run for ∼15 h at 25 w. The RNA product was visualized with UV shadowing, excised from the gel, and extracted by electroelution. The eluted RNA was spin concentrated, washed with ultra-pure sodium chloride, and exchanged into water using a 3K Amicon Ultra centrifugal filter. RNA purity was confirmed on a 10% analytical denaturing polyacrylamide gel and the concentration was quantified via UV-Vis absorbance. ^15^N/^13^C-adenosine (^15^N/^13^C-A) labeled NPSL2 was prepared as described above except that unlabeled rATP was replaced with ^15^N/^13^C-rATP in the transcription reaction.

### NMR data acquisition, processing, and analysis

All NMR spectra were acquired on either a Bruker 600 or 800 MHz Bruker ADVANCE NEO spectrometer, equipped with a 5 mm TCI cryogenic probe. Imino proton NMR data, for analysis of the NPSL2 secondary structure, were collected on 200 μM NPSL2 prepared in NMR buffer [50 mM potassium phosphate (pH = 7.5), 1 mM MgCl_2_] with varying concentrations of DMSO (v/v ratios) and 10% (v/v) D_2_O to a final volume of 300 μL in a 5 mm shigemi tube. Data were processed with TopSpin 4.1.4 and analyzed with MestreNova.^47^

NMR samples for STD experiments were prepared in NMR buffer in 5% v/v DMSO and 95% v/v D_2_O). Final concentrations were 10 μM NPSL2 and ∼ 200 μM ligand in 550 μL sample volumes. Reference ^1^H spectra for all ligands were collected under identical conditions without NPSL2. STD experiments were performed using a standard Bruker pulse sequence (stddiffesgp.3). The on-resonance irradiation frequency was set to 4.9 or 5.0 ppm, depending on the reference ligand ^1^H spectra obtained. Off-resonance irradiation was set to –40 ppm, where no RNA resonances are present. A total of 256 scans were acquired for each STD experiment, at 20 °C. Pre saturation of RNA resonances was achieved by an appropriate number of band-selective G4 Gaussian cascade pulses to give a saturation time of 2 s. Data was processed with TopSpin 4.1.4 and analyzed with MestReNova.^47^

NMR samples for HSQC experiments were prepared in NMR buffer containing 10% v/v D_2_O and 5% v/v DMSO. Final concentrations were 100 μM ^15^N/^13^C-A labeled NPSL2 to up to 1 mM ligand (depending on the solubility of the ligand) in a total volume of 150 μL (3 mm tube). The temperature was set to 15 °C to match our previously acquired and assigned HSQC spectra (BMRB ID: 51350).^21^ Data were processed with NMRFx^48^ and analyzed with NMRViewJ.^49^ All resonance assignments for ligands were predicted using ChemDraw.

### CD thermal denaturation and data analysis

Thermal denaturation experiments were performed on a JASCO J-815 CD spectrometer. The heating rate was set to 1 °C per min and ramped from 5 °C to 95 °C with detection at 260 nm. Data points were collected every 0.5 °C. Samples were prepared by mixing 10 μM NPSL2 with 200 μM ligand [in 50 mM potassium phosphate buffer (pH = 7.5), 1 mM MgCl_2_, 5% v/v DMSO]. The single-transition melting profiles were analyzed by a two-state model.^50^

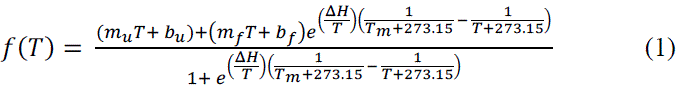

Where m_u_ and m_f_ are the slopes of the unfolded and folded baselines, and b_u_ and b_f_ are the y-intercepts of the unfolded and folded baselines, respectively. Δ*H* (Kcal/mol) and *T_m_* (°C) are the enthalpy of folding and melting temperature, respectively. Δ*S* was determined using the following equation.

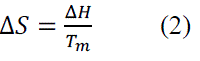

Experiments were performed in triplicate. The average and standard deviation are reported in **Table S2**.

## DATA AVAILABILITY

The datasets generated during and/or analyzed during the current study are available from the corresponding author on reasonable request.

## ACKNOWLEDGEMENTS

This work was supported by the National Institutes of Health [R35 GM138279 to S.C.K.]. Research reported in this publication was supported by the University of Michigan BioNMR Core Facility (U-M BioNMR). U-M BioNMR Core is grateful for support from U-M including the College of Literature, Sciences and Arts, Life Sciences Institute, College of Pharmacy, and the Medical School along with the U-M Biosciences Initiative. We would like to thank Prof. Charles L. Brooks III for giving us access to the Gollum computing cluster where all modelling related work was performed. We thank Debashish Sahu and Minli Xing for technical assistance with NMR data acquisition.

## AUTHOR CONTRIBUTIONS

S.C.K. conceived project. G.A. conducted all computations and experiments. G.A. and S.C.K. analyzed the results. G.A. and S.C.K wrote the manuscript.

## ADDITIONAL INFORMATION

The authors declare no conflict of interest.

## Notes

### Competing Interest Statement

The authors have declared no competing interest.

