## Supplementary Data for "*In silico* discovery of compounds targeting NPSL2: a regulatory element in the human oncomiR-1 primary microRNA"

**Table S1**. Docking score free energies across 3 different conformers based on predicted binding pocket. All compounds are shown by their ZINC IDs and abbreviated three number codes used in later discussions

| Compound ID | Apical loop score (Kcal/mol) | Internal loop score (Kcal/mol) | Dual loop score (Kcal/mol) |
|---|---|---|---|
| ZINC000021659043 (Zn 043) | -22.6 | -26.9 | -20.8 |
| ZINC000002059079 (Zn 079) | -16.9 | -27.2 | -19.0 |
| ZINC000002059091 (Zn 091) | -23.0 | -25.2 | -18.8 |
| ZINC000021659126 (Zn 126) | -16.3 | -30.3 | -22.9 |
| ZINC000013688273 (Zn 273) | -12.9 | -21.8 | -21.1 |
| ZINC000002312347 (Zn 347) | -16.4 | -22.8 | -19.5 |
| ZINC000006148504 (Zn 504) | -16.4 | -31.4 | -15.9 |
| ZINC000016755513 (Zn 513) | -18.5 | -20.1 | -22.0 |
| ZINC000000902588 (Zn 588) | -18.8 | -26.9 | -20.6 |
| ZINC000096221591 (Zn 591) | -23.4 | -24.2 | -18.9 |
| ZINC000000902668 (Zn 668) | -17.9 | -28.1 | -18.9 |
| ZINC000006878699 (Zn 699) | -19.8 | -25.5 | -19.6 |
| ZINC000096222711 (Zn 711) | -21.7 | -21.3 | -17.8 |
| ZINC000033544735 (Zn 735) | -20.5 | -22.2 | -24.3 |
| ZINC000097971918 (Zn 918) | -17.0 | -21.1 | -20.7 |
| ZINC000103558981 (Zn 981) | -15.2 | -31.8 | -16.4 |

**Table S2**. Thermodynamic properties of NPSL2-ligand complexes

| ZINC ID | $T_m$[a] (℃) | ΔH[a] (kcal/mol) | ΔS[a] [cal/(mol*K)] | $\Delta T_m$ (℃) | ΔΔH (kcal/mol) | ΔΔS [cal/(mol*K)] |
|---|---|---|---|---|---|---|
| DMSO | 68.8 ± 0.1 | -96.6 ± 6.0 | -282.7 ± 17.4 | - | - | - |
| Zn 043 | 68.8 ± 0.3 | -95.6 ± 3.7 | -279.7 ± 10.9 | 0.0 | 1.0 | 3.0 |
| Zn 126 | 68.9 ± 0.2 | -89.4 ± 6.6 | -261.4 ± 19.3 | 0.1 | 7.3 | 21.3 |
| Zn 588 | 68.9 ± 0.2 | -109.3 ± 6.5 | -319.7 ± 19.1 | 0.1 | -12.7 | -37.0 |
| Zn 735 | 68.7 ± 0.3 | -91.1 ± 6.6 | -266.5 ± 19.5 | -0.1 | 5.5 | 16.1 |
| Zn 699 | 68.5 ± 0.3 | -104.3 ± 9.2 | -305.3 ± 27.1 | -0.3 | -7.6 | -22.6 |
| Zn 504 | 68.9 ± 0.3 | -93.3 ± 3.9 | -272.8 ± 11.4 | 0.1 | 3.3 | 9.8 |
| Zn 591 | 69.1 ± 0.3 | -75.6 ± 4.4 | -220.9 ± 12.9 | 0.3 | 21.0 | 61.7 |
| Zn 668 | 69.2 ± 0.4 | -83.8 ± 8.0 | -245.0 ± 23.4 | 0.4 | 12.7 | 37.7 |

[a]Average values and standard deviations are reported from triplicate experiments

**Table S3**. Compounds with number of hydrogen bonding donors and acceptors

| Compound ID | H-bond donors | H-bond acceptors |
|---|---|---|
| Zn 126 | 3 | 8 |
| Zn 079 | 1 | 4 |
| Zn 091 | 1 | 4 |
| Zn 699 | 3 | 13 |
| Zn 588 | 4 | 11 |
| Zn 668 | 3 | 11 |
| Zn 711 | 3 | 7 |
| Zn 918 | 5 | 10 |
| Zn 591 | 5 | 9 |
| Zn 981 | 1 | 7 |
| Zn 504 | 1 | 5 |
| Zn 273 | 2 | 9 |
| Zn 043 | 3 | 8 |
| Zn 513 | 3 | 8 |
| Zn 347 | 1 | 6 |
| Zn 735 | 3 | 9 |

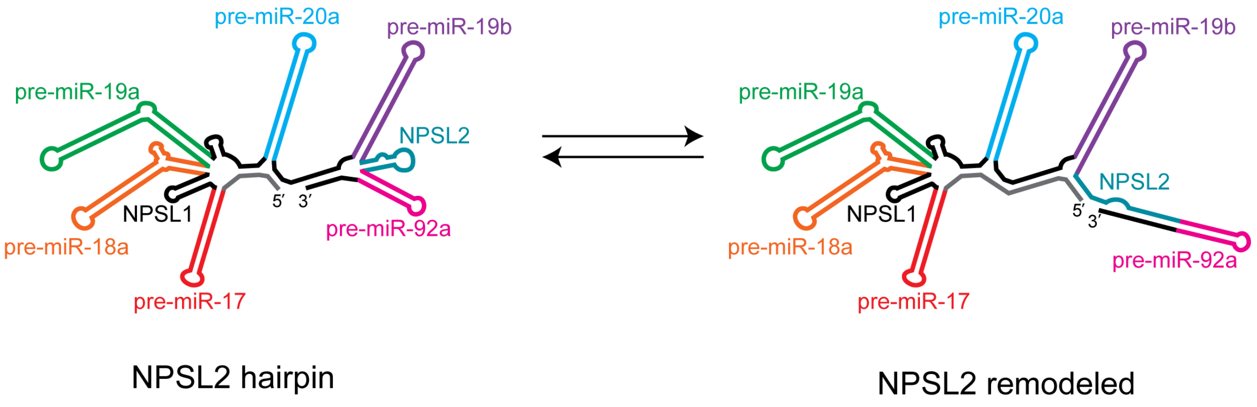


**Figure S1.** Predicted conformational rearrangement within the 3′-region of oncomiR-1.

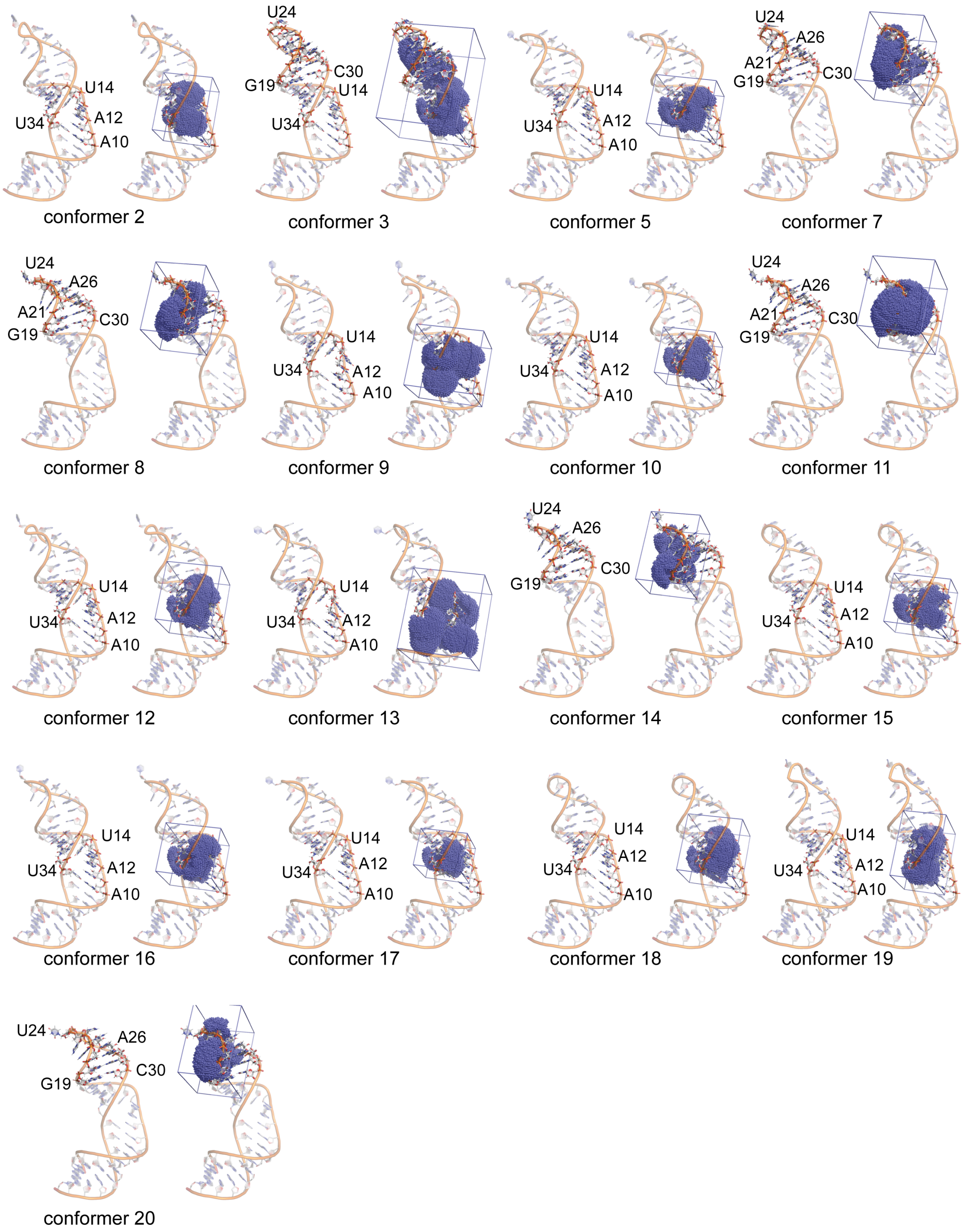


**Figure S2**. 3D structure of NPSL2 conformers showing the apical and internal loops (left) with their predicted binding volumes (in blue dots and a square box, right) determined using RNACavityMiner.[1]

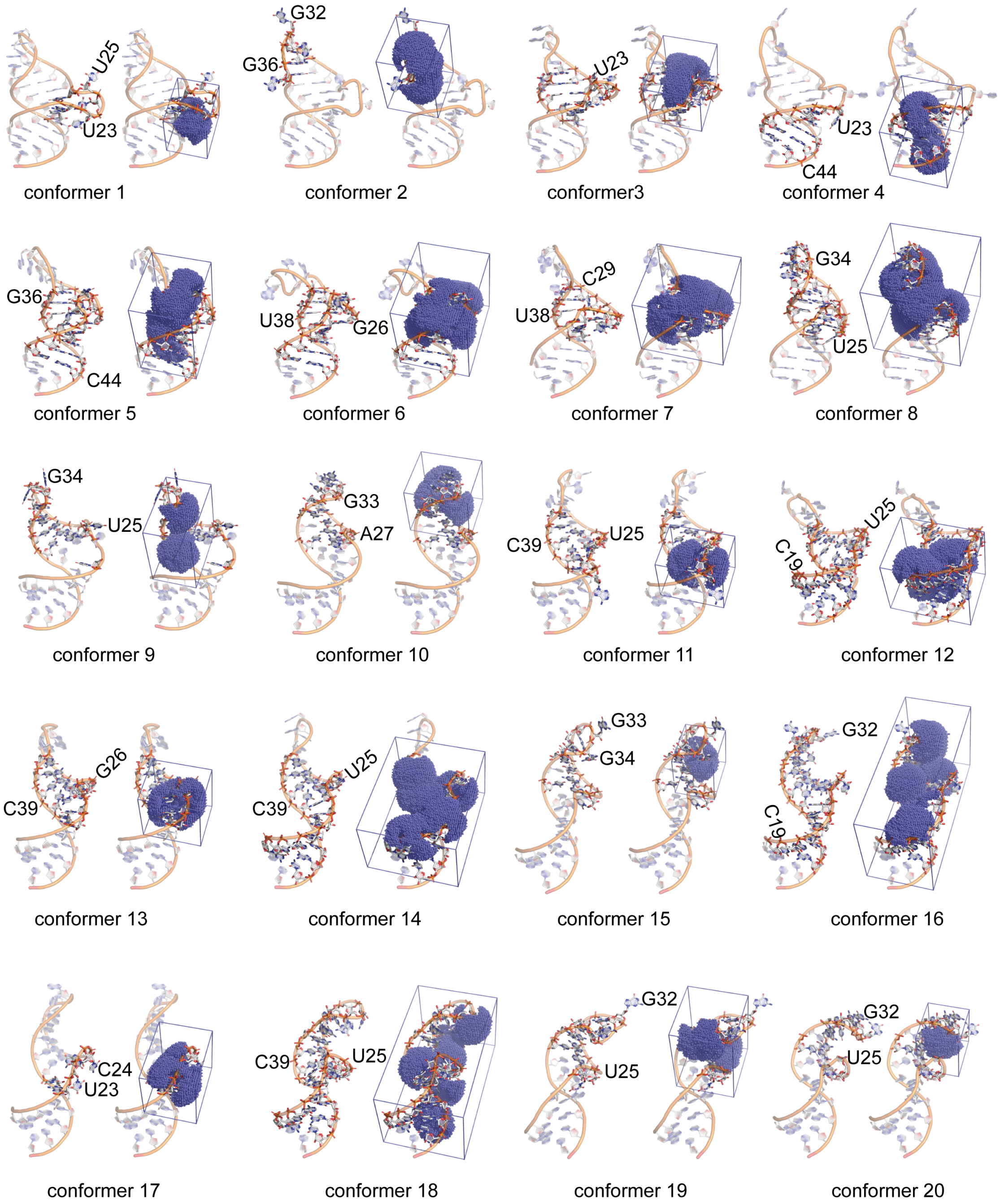


**Figure S3**. 3D structure of TAR RNA conformers showing the apical and internal loops (left) with their predicted binding volumes (in blue dots and a square box, right) determined using RNACavityMiner.[1]

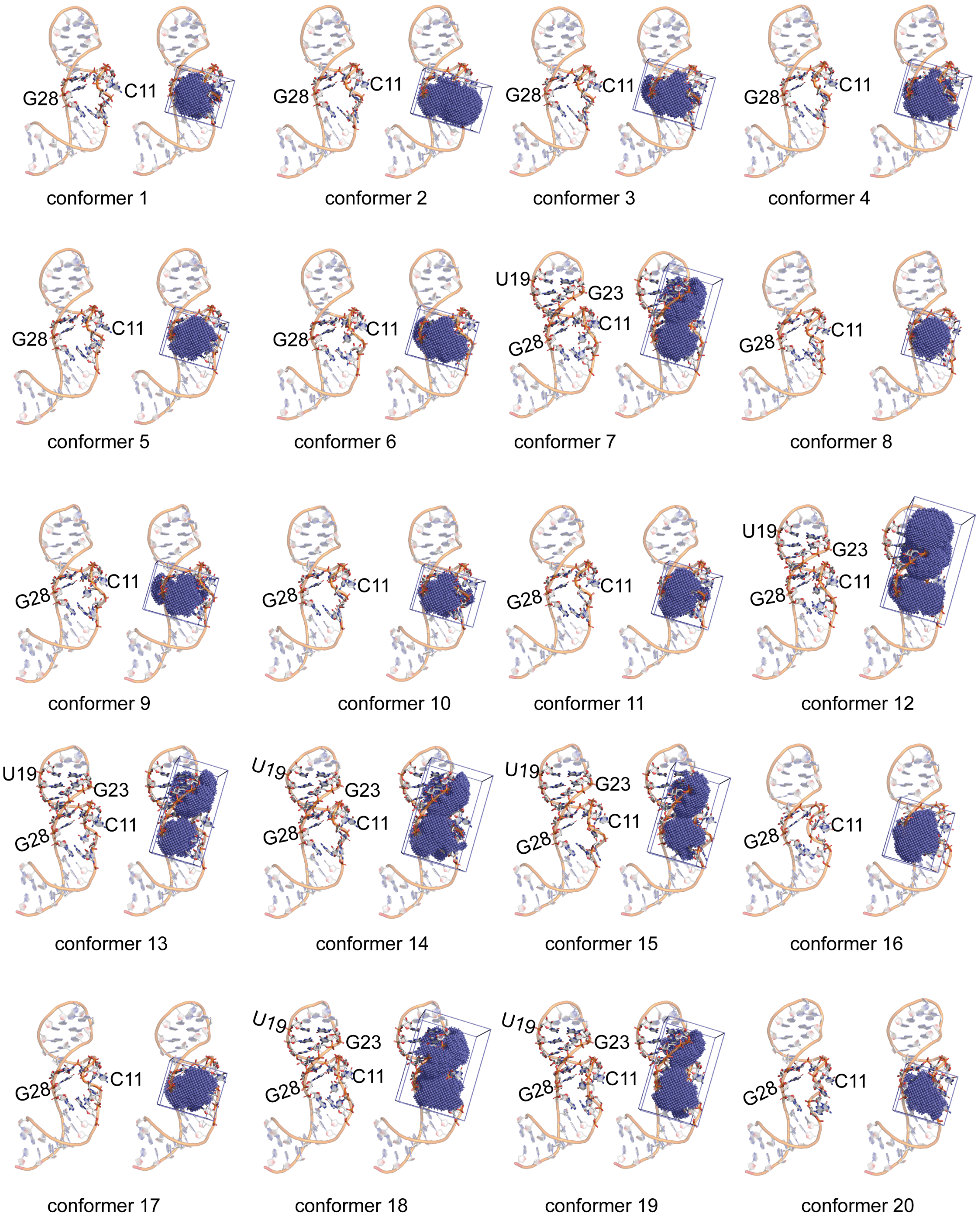


**Figure S4**. 3D structures of telomerase RNA conformers showing the apical and internal loops (left) with their predicted binding volumes (in blue dots and a square box, right) determined using RNACavityMiner.[1]

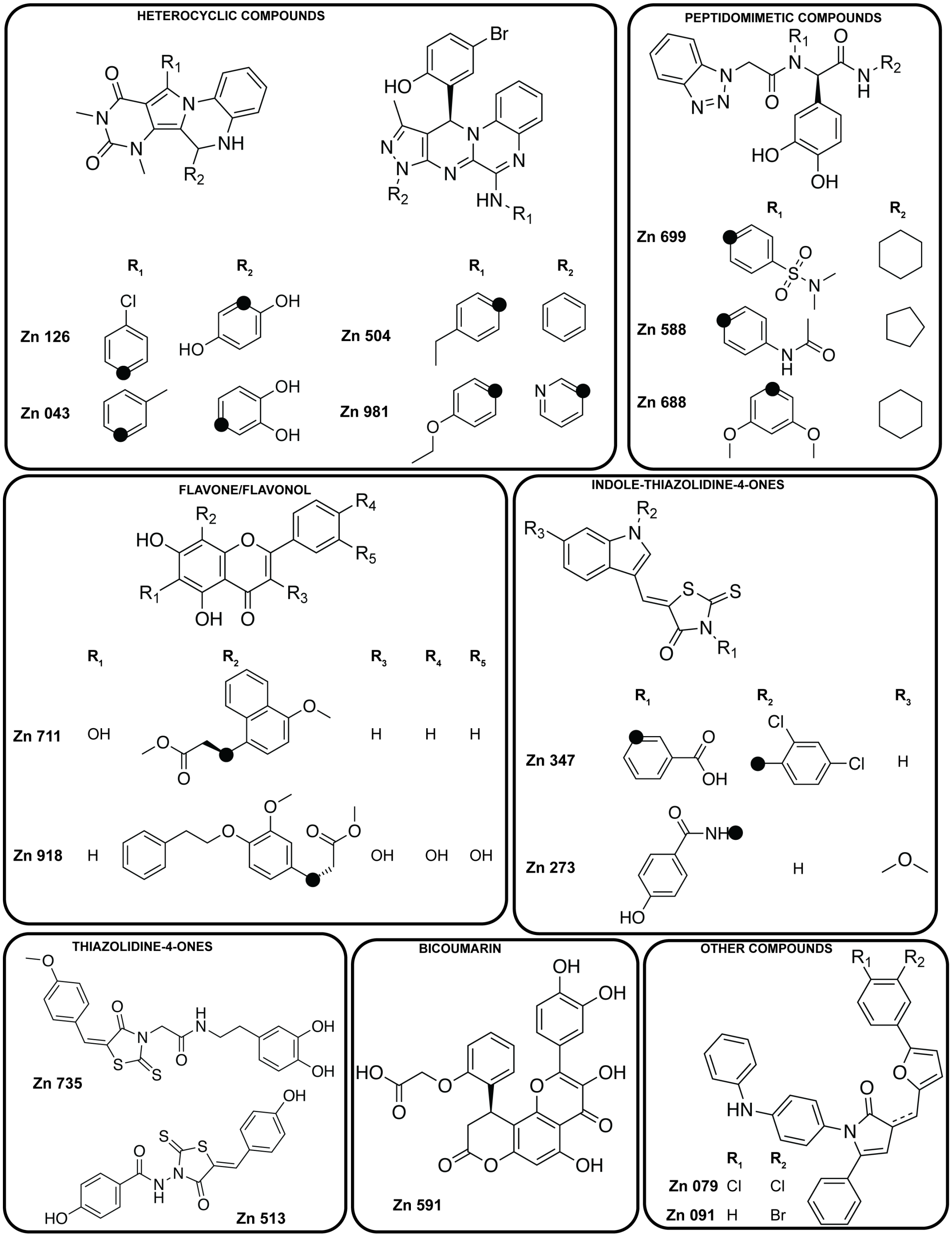


**Figure S5**. Chemical structures of purchased compounds grouped based on shared scaffolds/characteristics. Black circles indicate the point of connection to the scaffold.

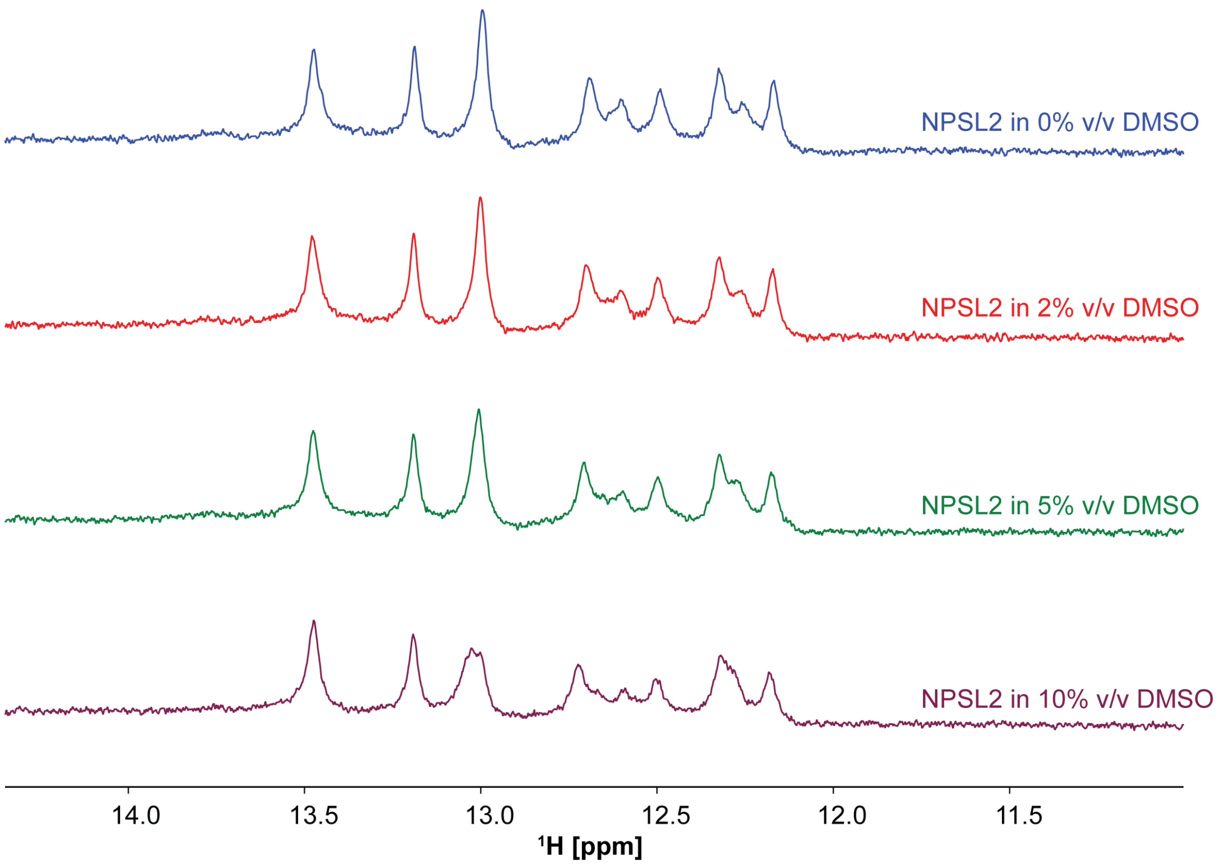


**Figure S6**. Imino proton spectra of NPSL2 prepared in buffer containing different amounts of DMSO. The NMR spectra were recorded on NPSL2 RNA (~ 300 μM) in buffer containing 50 mM $KH_2PO_4$, pH = 7.5, 1 mM $MgCl_2$, 10% $D_2O$, and DMSO ranging from 0 – 10 % (v/v).

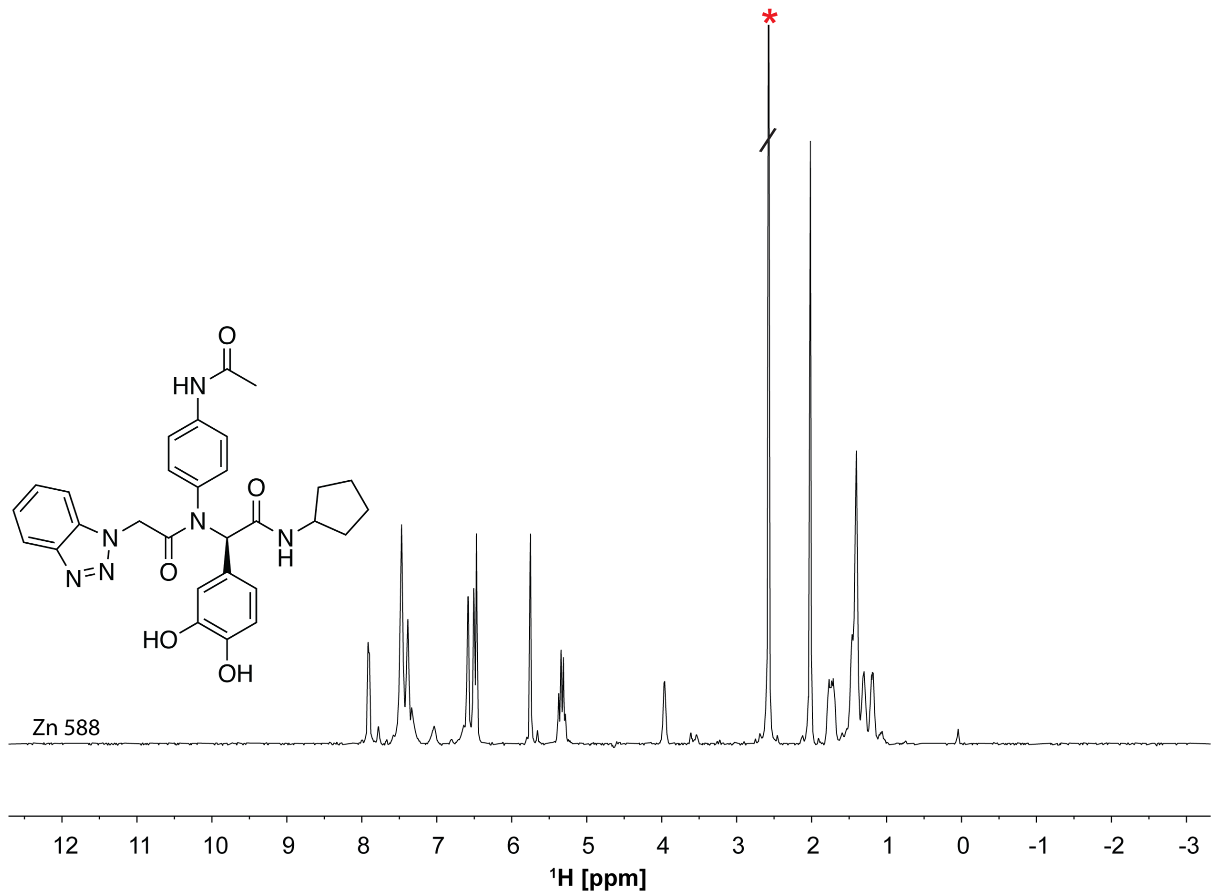


**Figure S7.** Chemical structure and ¹H spectrum of Zn 588. The NMR spectrum was recorded at ~ 200 µM concentration, 50 mM $KH_2PO_4$, pH = 7.5, 50 mM KCl, 1 mM $MgCl_2$, 95% $D_2O$ and 5% DMSO on a 600 MHz Bruker spectrometer equipped with a cryoprobe. The red asterisk indicates the DMSO peak.

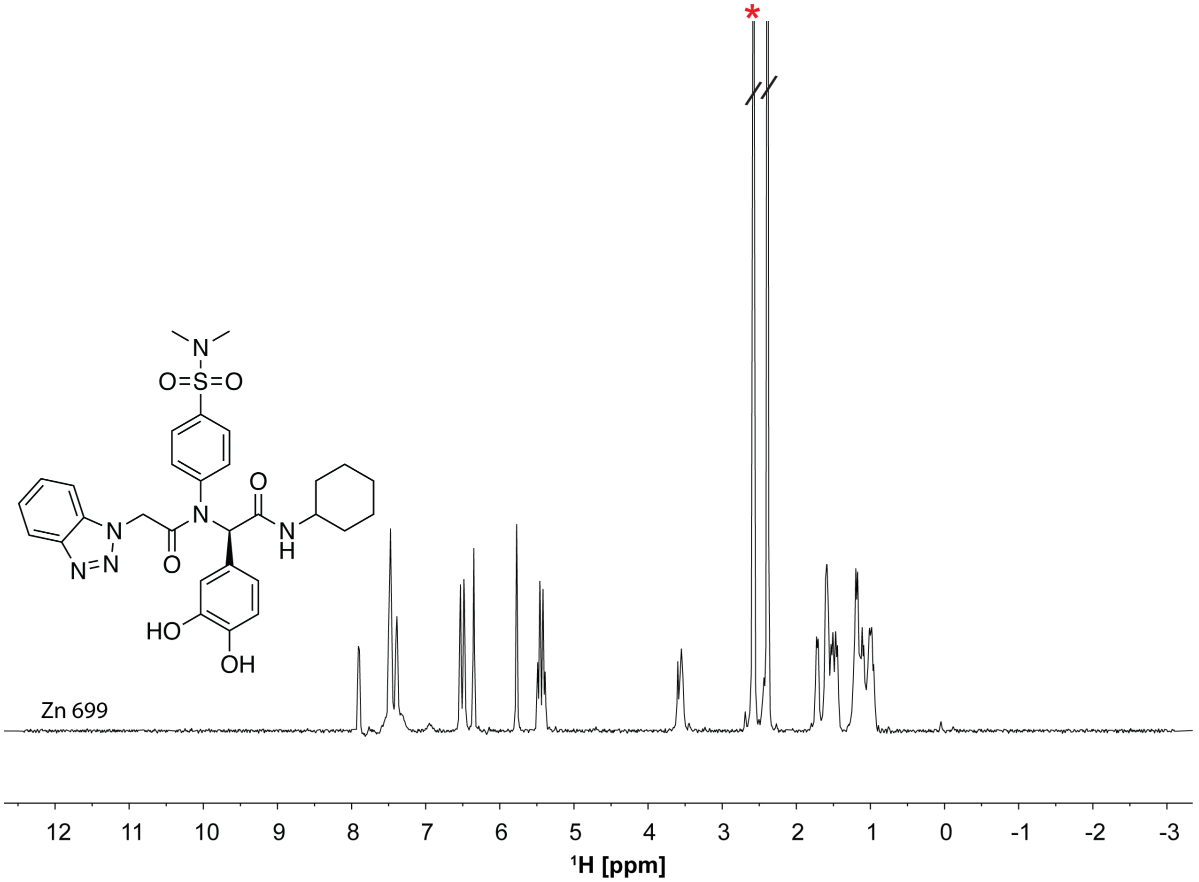


**Figure S8.** Chemical structure and $^{1}$H spectrum of Zn 699. The NMR spectra was recorded at ~ 200 µM concentration, 50 mM $KH_2PO_4$, pH = 7.5, 50 mM KCl, 1 mM $MgCl_2$, 95% $D_2O$ and 5% DMSO on a 600 MHz Bruker spectrometer equipped with a cryoprobe. The red asterisk indicates the DMSO peak.

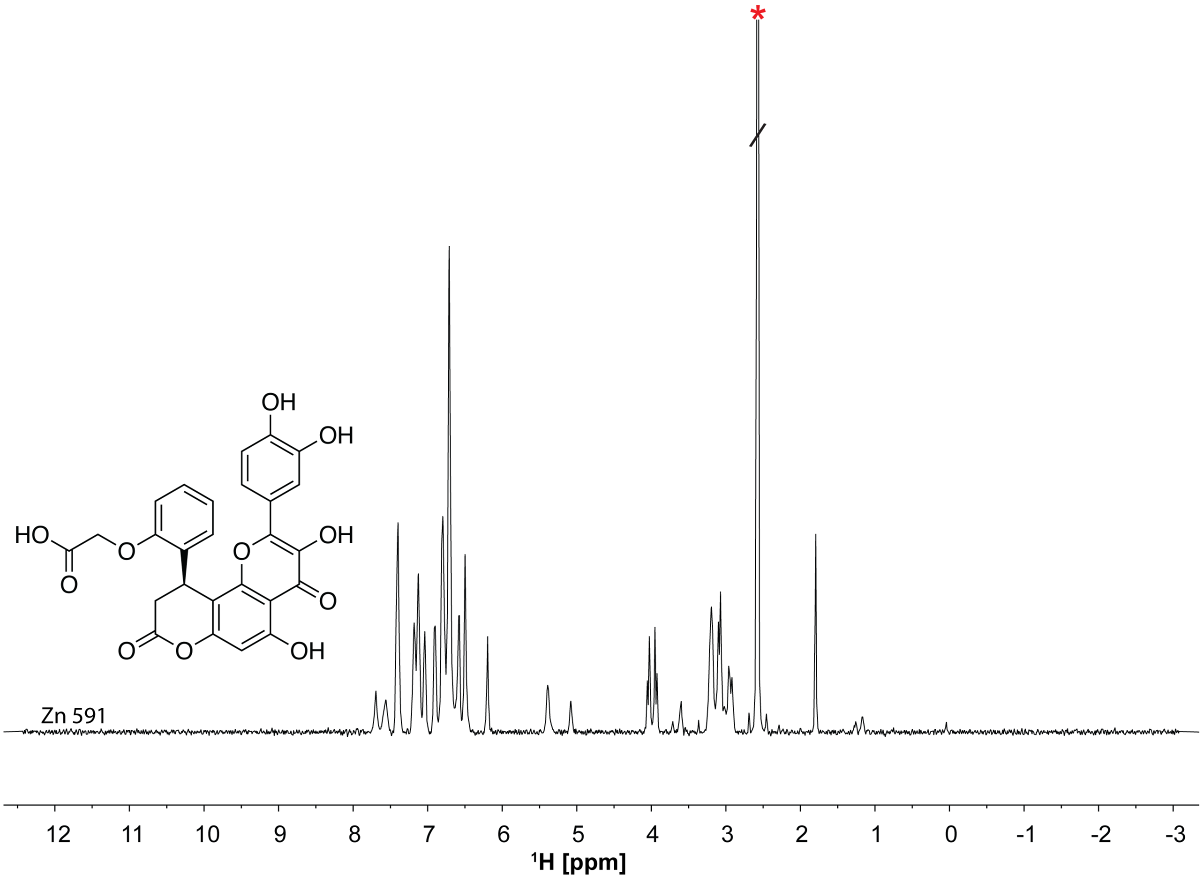


**Figure S9.** Chemical structure and $^{1}H$ spectrum of Zn 591. The NMR spectra was recorded at ~ 200 μM concentration, 50 mM $KH_2PO_4$, pH = 7.5, 50 mM KCl, 1 mM $MgCl_2$, 95% $D_2O$ and 5% DMSO on a 600 MHz Bruker spectrometer equipped with a cryoprobe. The red asterisk indicates the DMSO peak.

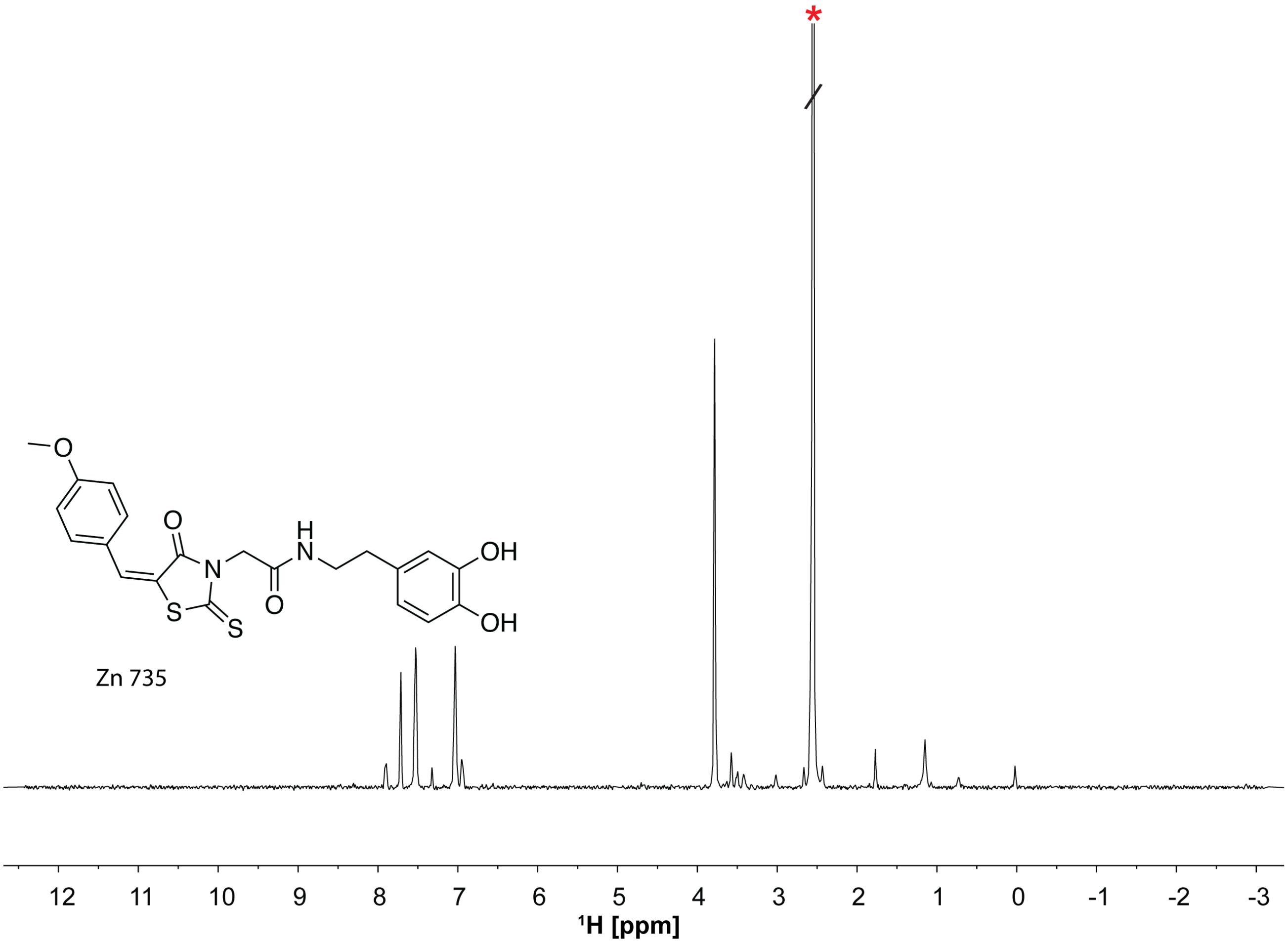


**Figure S10: [1]H spectrum of Zn 735 compound showing 2D structure.** The NMR spectra was recorded at ~ 200 µM concentration, 50 mM $KH_2PO_4$, pH = 7.5, 50 mM KCl, 1 mM $MgCl_2$, 95% $D_2O$ and 5% DMSO on a 600 MHz Bruker spectrometer equipped with a cryoprobe. The red asterisk indicates the DMSO peak.

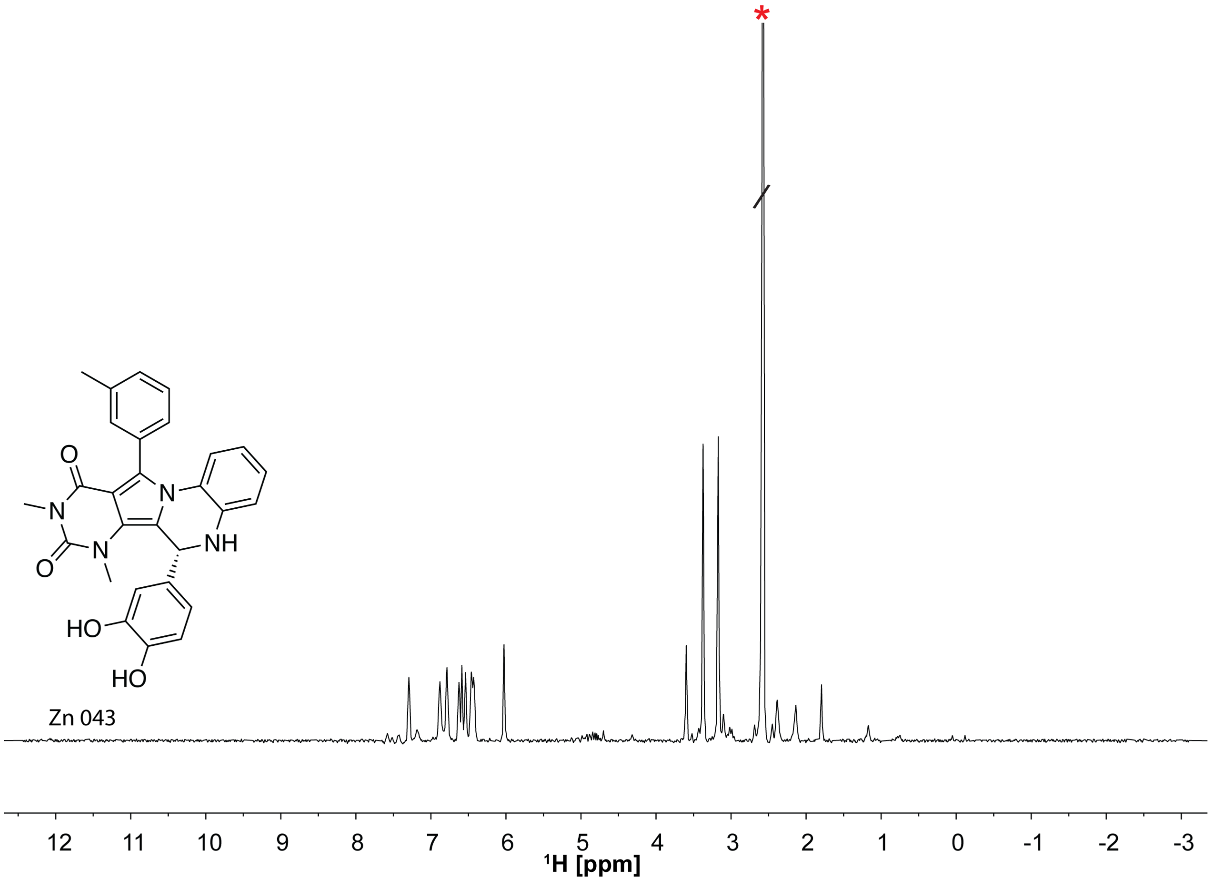


**Figure S11.** Chemical structure and $^{1}H$ spectrum of Zn 043. The NMR spectra was recorded at ~ 200 µM concentration, 50 mM $KH_2PO_4$, pH = 7.5, 50 mM KCl, 1 mM $MgCl_2$, 95% $D_2O$ and 5% DMSO on a 600 MHz Bruker spectrometer equipped with a cryoprobe. The red asterisk indicates the DMSO peak.

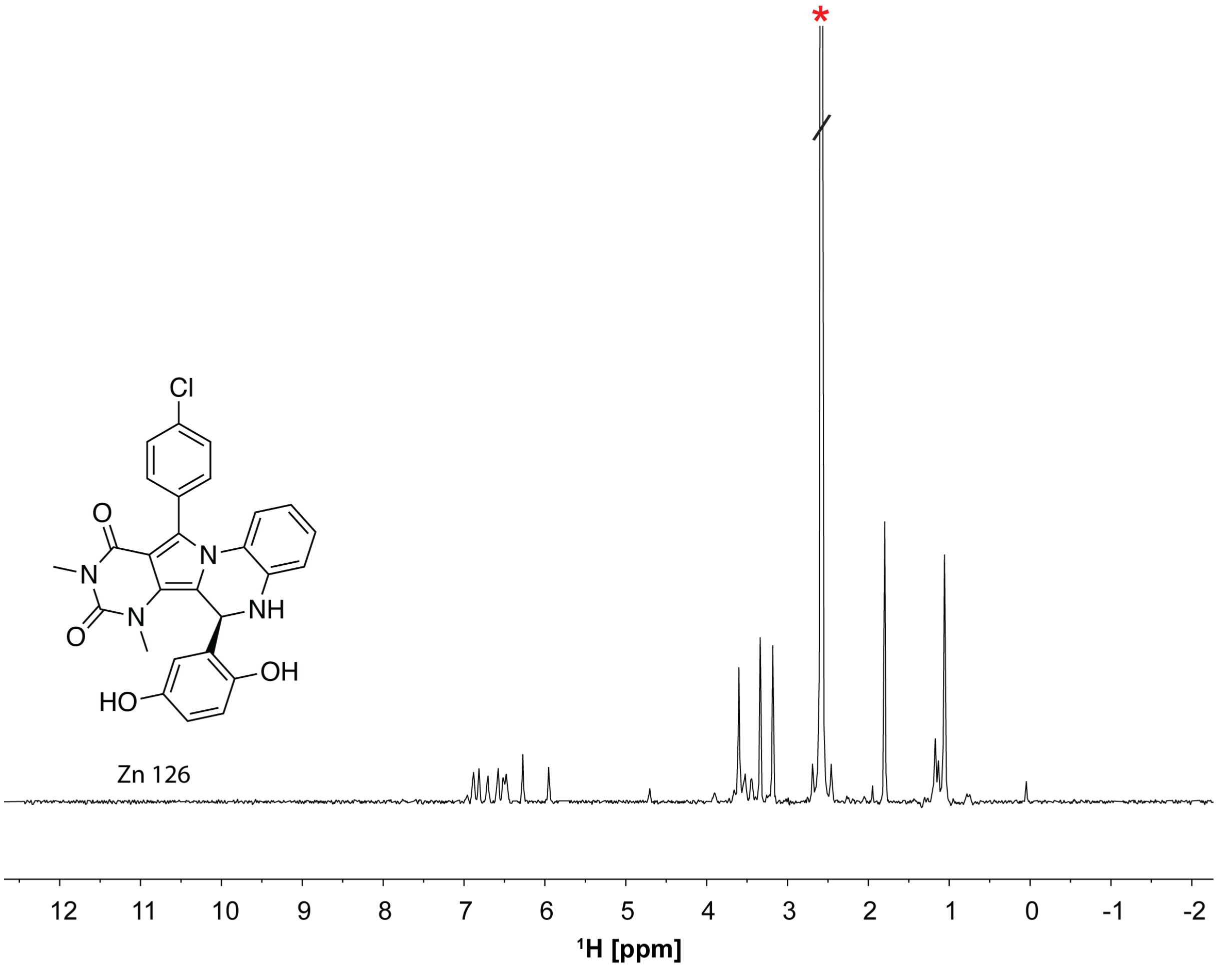


**Figure S12.** Chemical structure and ¹H spectrum of Zn 126. The NMR spectra was recorded at ~ 200 μM concentration, 50 mM $KH_2PO_4$, pH = 7.5, 50 mM KCl, 1 mM $MgCl_2$, 95% $D_2O$ and 5% DMSO on a 600 MHz Bruker spectrometer equipped with a cryoprobe. The red asterisk indicates the DMSO peak.

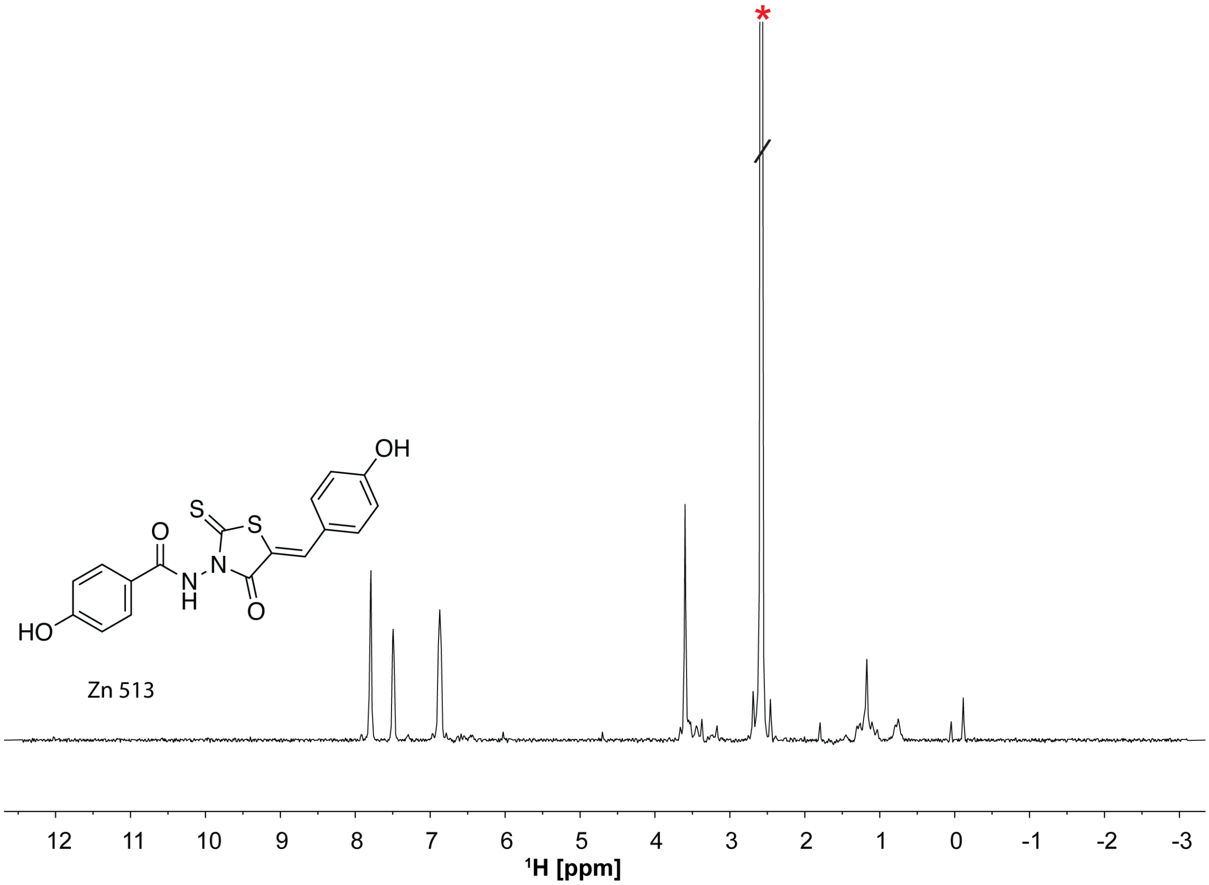


**Figure S13.** Chemical structure and [1]H spectrum of Zn 513. The NMR spectra was recorded at ~ 200 μM concentration, 50 mM $KH_2PO_4$, pH = 7.5, 50 mM KCl, 1 mM $MgCl_2$, 95% $D_2O$ and 5% DMSO on a 600 MHz Bruker spectrometer equipped with a cryoprobe. The red asterisk indicates the DMSO peak.

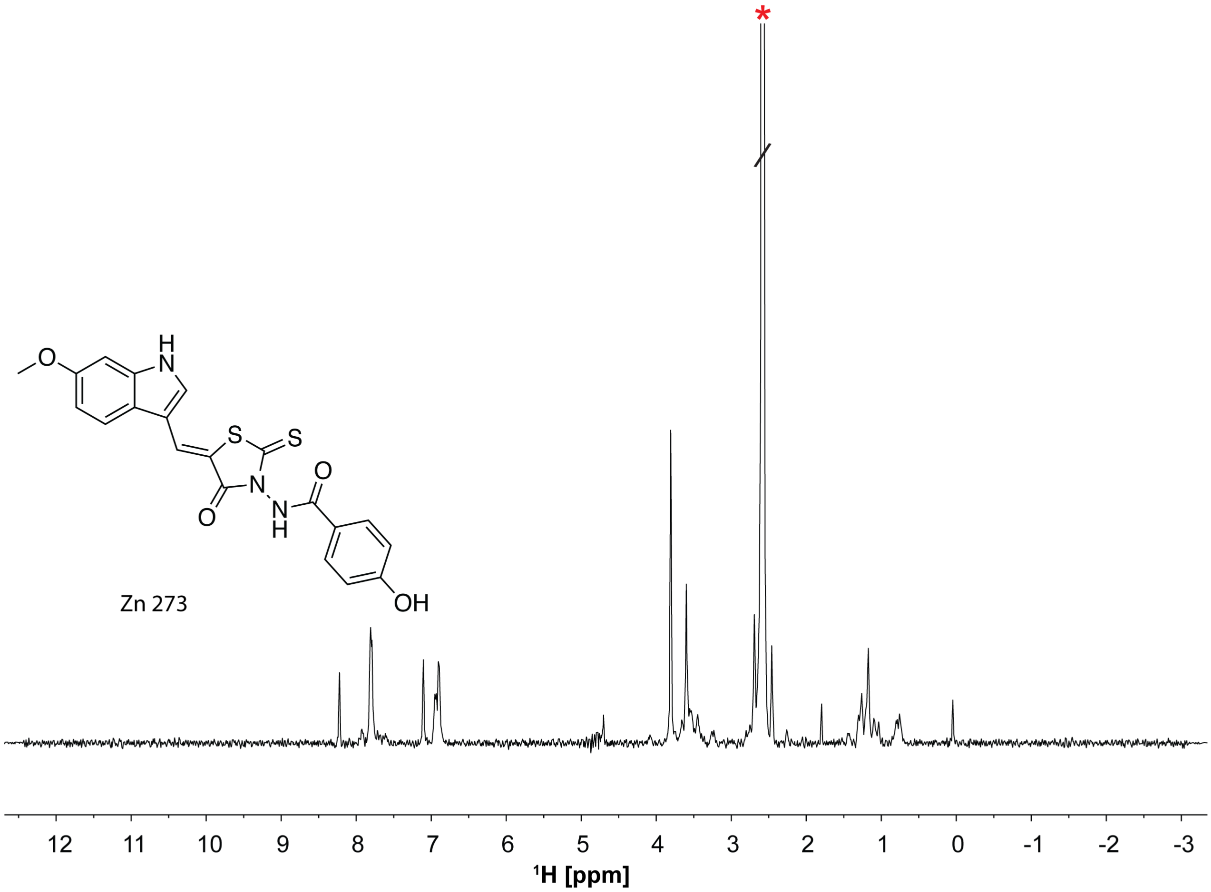


**Figure S14.** Chemical structure and [1]H spectrum of Zn 273. The NMR spectra was recorded at ~ 200 µM concentration, 50 mM $KH_2PO_4$, pH = 7.5, 50 mM KCl, 1 mM $MgCl_2$, 95% $D_2O$ and 5% DMSO on a 600 MHz Bruker spectrometer equipped with a cryoprobe. The red asterisk indicates the DMSO peak.

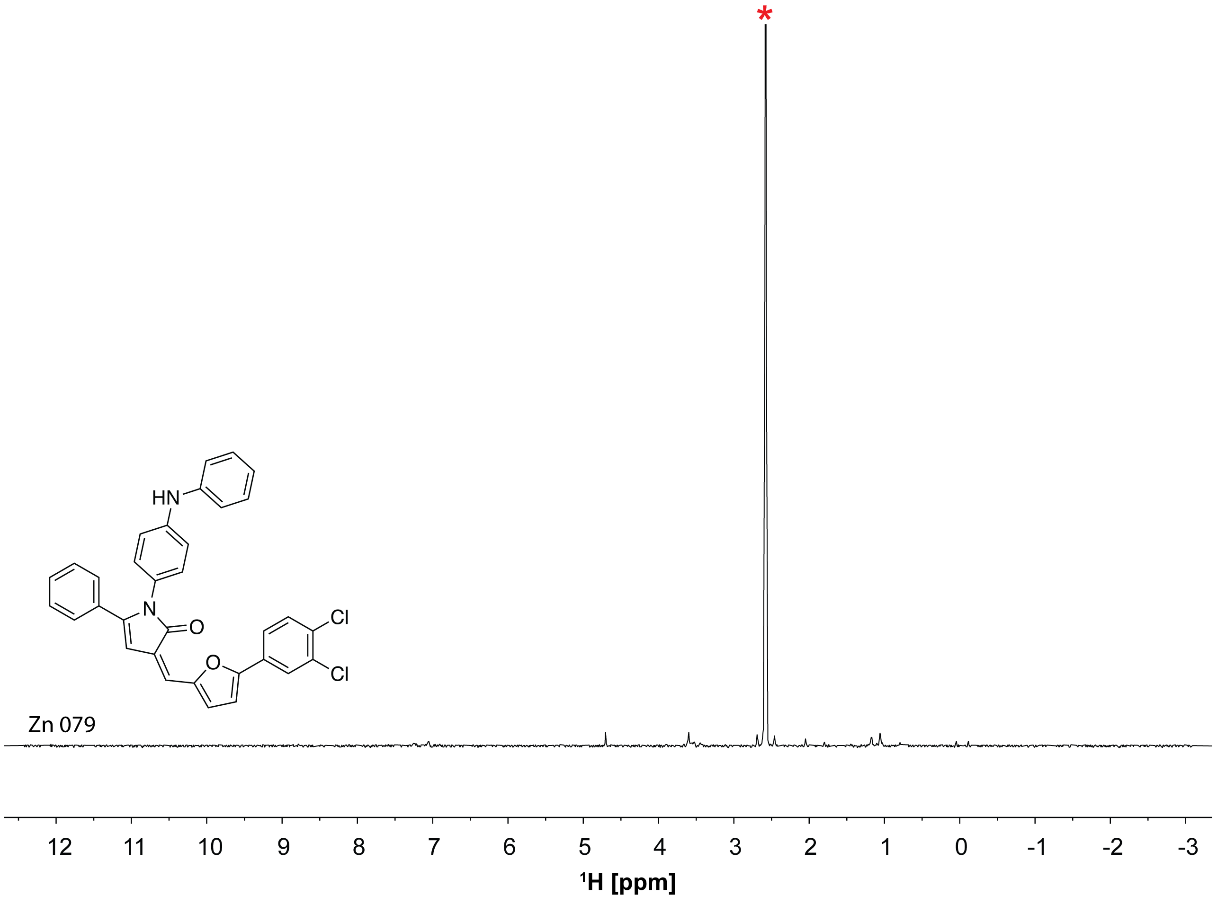


**Figure S15.** Chemical structure and $^{1}H$ spectrum of Zn 079. The NMR spectrum was recorded on Zn 079 solubilized in buffer containing 50 mM $KH_2PO_4$, pH = 7.5, 50 mM KCl, 1 mM $MgCl_2$, 95% $D_2O$ and 5% DMSO on a 600 MHz Bruker spectrometer equipped with a cryoprobe. The red asterisk indicates the DMSO peak.

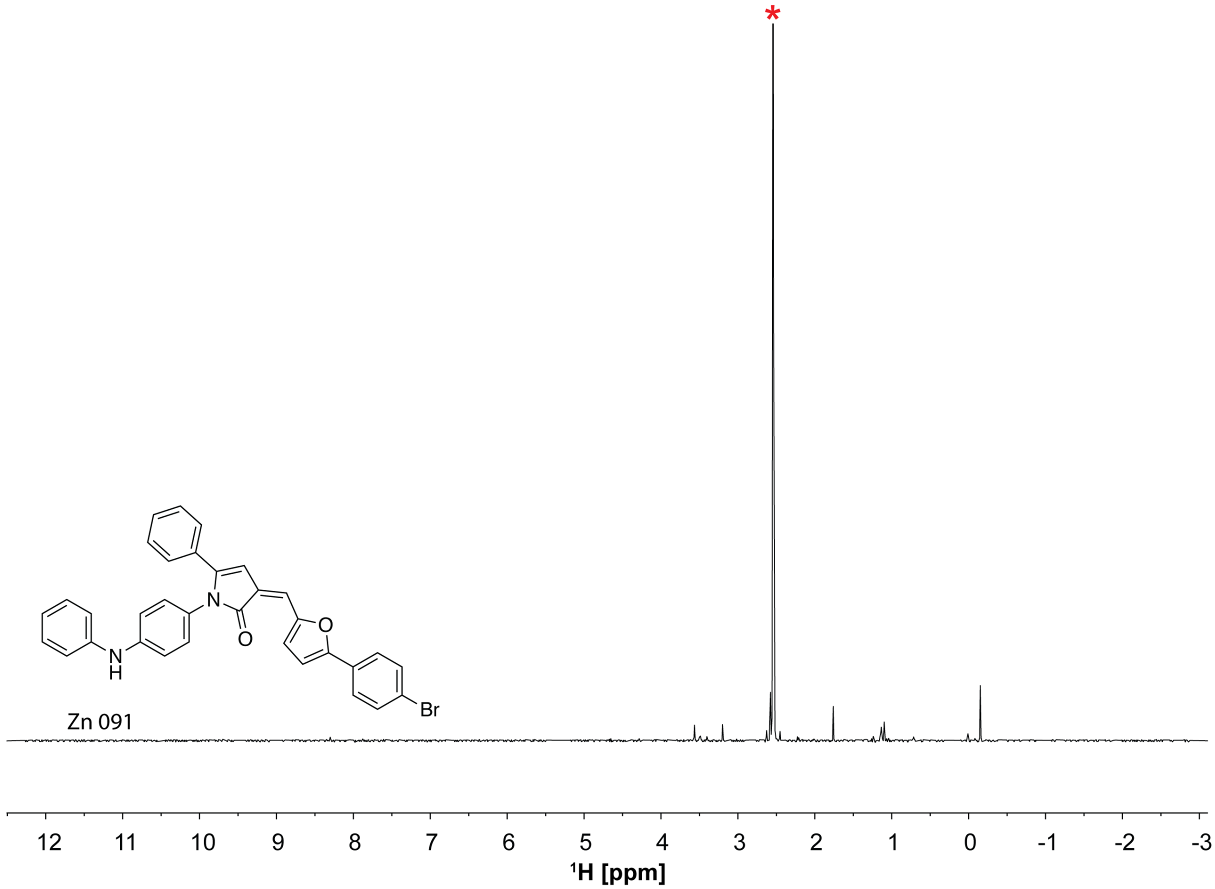


**Figure S16.** Chemical structure and $^1$H spectrum of Zn 091. The NMR spectrum was recorded on Zn 091 solubilized in buffer containing 50 mM $KH_2PO_4$, pH = 7.5, 50 mM KCl, 1 mM $MgCl_2$, 95% $D_2O$ and 5% DMSO on a 600 MHz Bruker spectrometer equipped with a cryoprobe. The red asterisk indicates the DMSO peak.

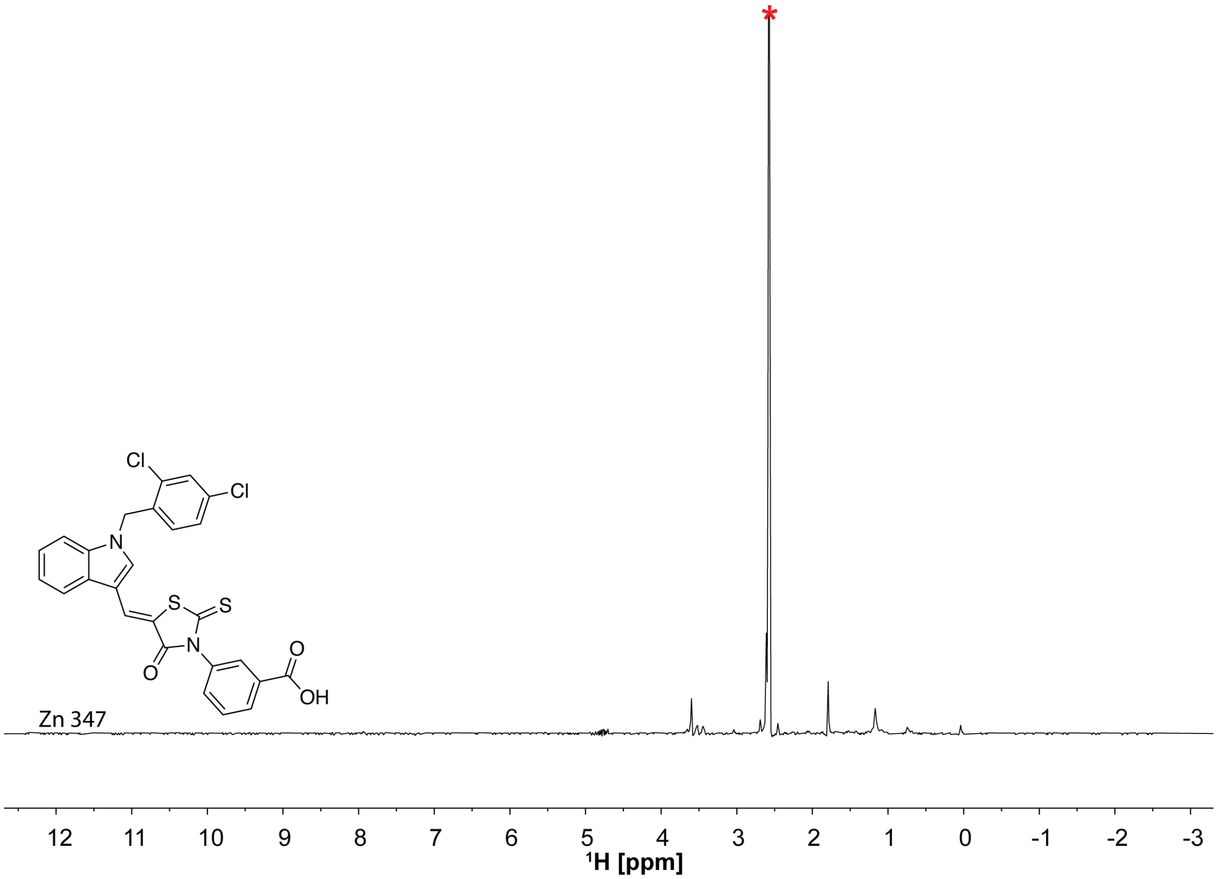


**Figure S17.** Chemical structure and $^{1}H$ spectrum of Zn 347. The NMR spectrum was recorded on Zn 347 solubilized in buffer containing 50 mM $KH_2PO_4$, pH = 7.5, 50 mM KCl, 1 mM $MgCl_2$, 95% $D_2O$ and 5% DMSO on a 600 MHz Bruker spectrometer equipped with a cryoprobe. The red asterisk indicates the DMSO peak.

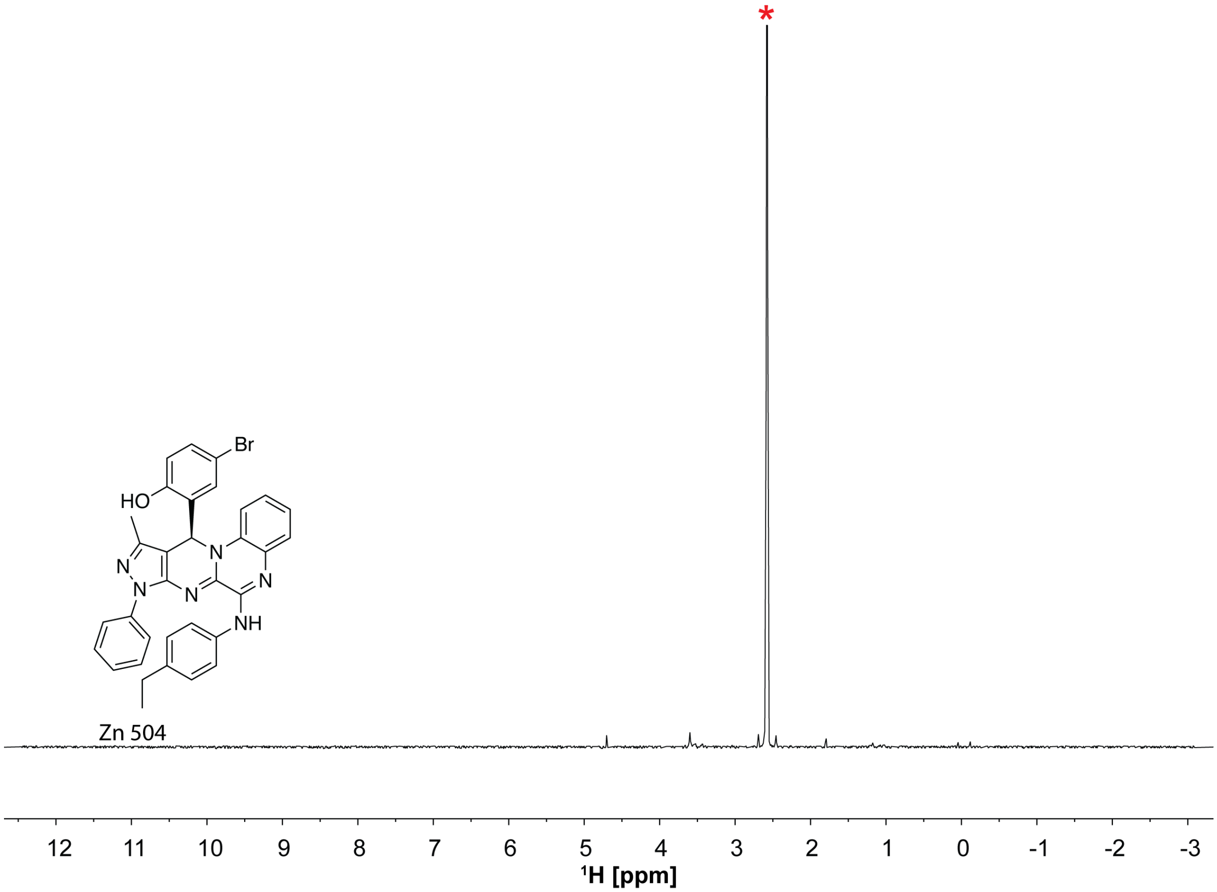


**Figure S18.** Chemical structure and $^1$H spectrum of Zn 504. The NMR spectrum was recorded on Zn 504 solubilized in buffer containing 50 mM $KH_2PO_4$, pH = 7.5, 50 mM KCl, 1 mM $MgCl_2$, 95% $D_2O$ and 5% DMSO on a 600 MHz Bruker spectrometer equipped with a cryoprobe. The red asterisk indicates the DMSO peak.

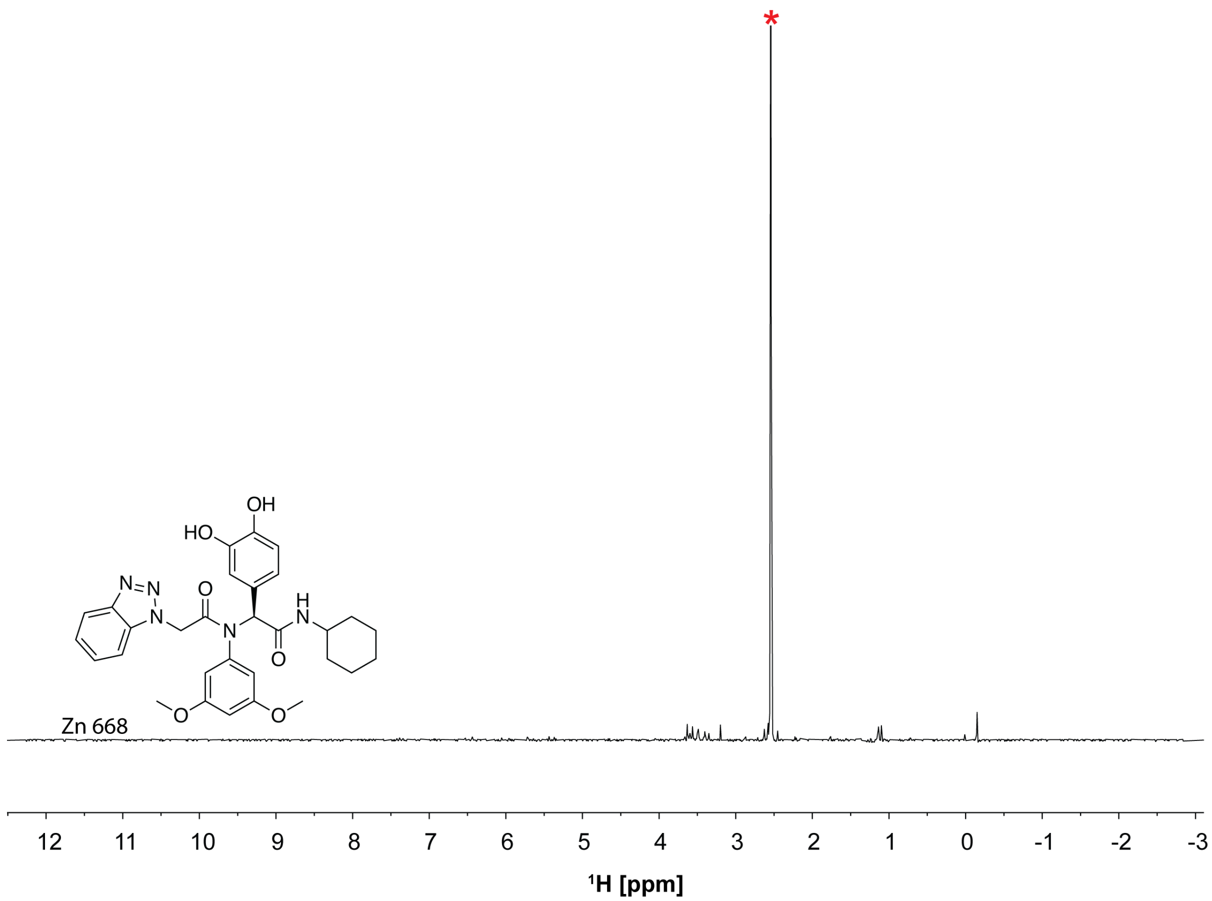


**Figure S19.** Chemical structure and $^1$H spectrum of Zn 668. The NMR spectrum was recorded on Zn 668 solubilized in buffer containing 50 mM $KH_2PO_4$, pH = 7.5, 50 mM KCl, 1 mM $MgCl_2$, 95% $D_2O$ and 5% DMSO on a 600 MHz Bruker spectrometer equipped with a cryoprobe. The red asterisk indicates the DMSO peak.

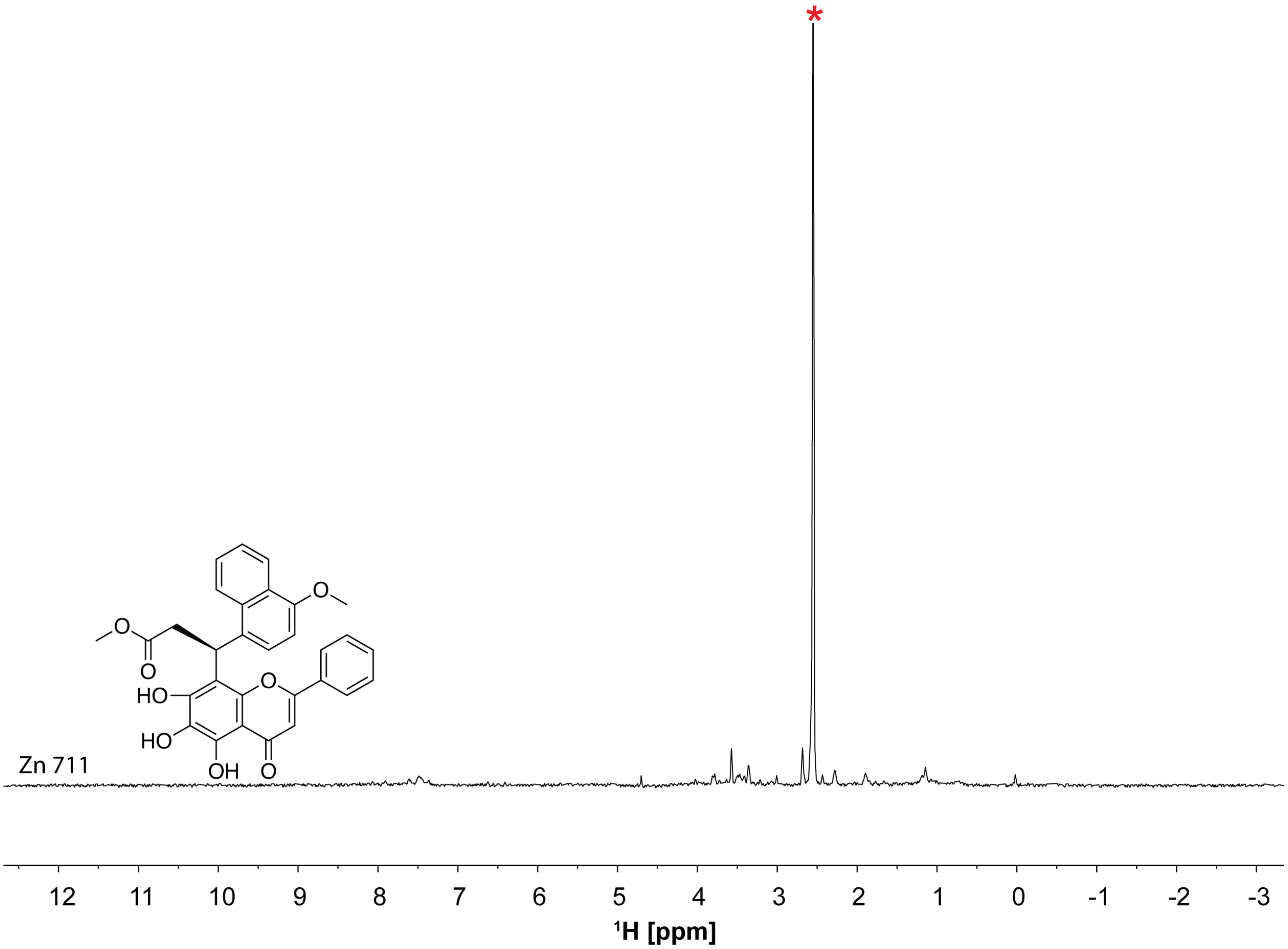


**Figure S20.** Chemical structure and [1]H spectra of Zn 711. The NMR spectrum was recorded on Zn 711 solubilized in buffer containing 50 mM $KH_2PO_4$, pH = 7.5, 50 mM KCl, 1 mM $MgCl_2$, 95% $D_2O$ and 5% DMSO on a 600 MHz Bruker spectrometer equipped with a cryoprobe. The red asterisk indicates the DMSO peak.

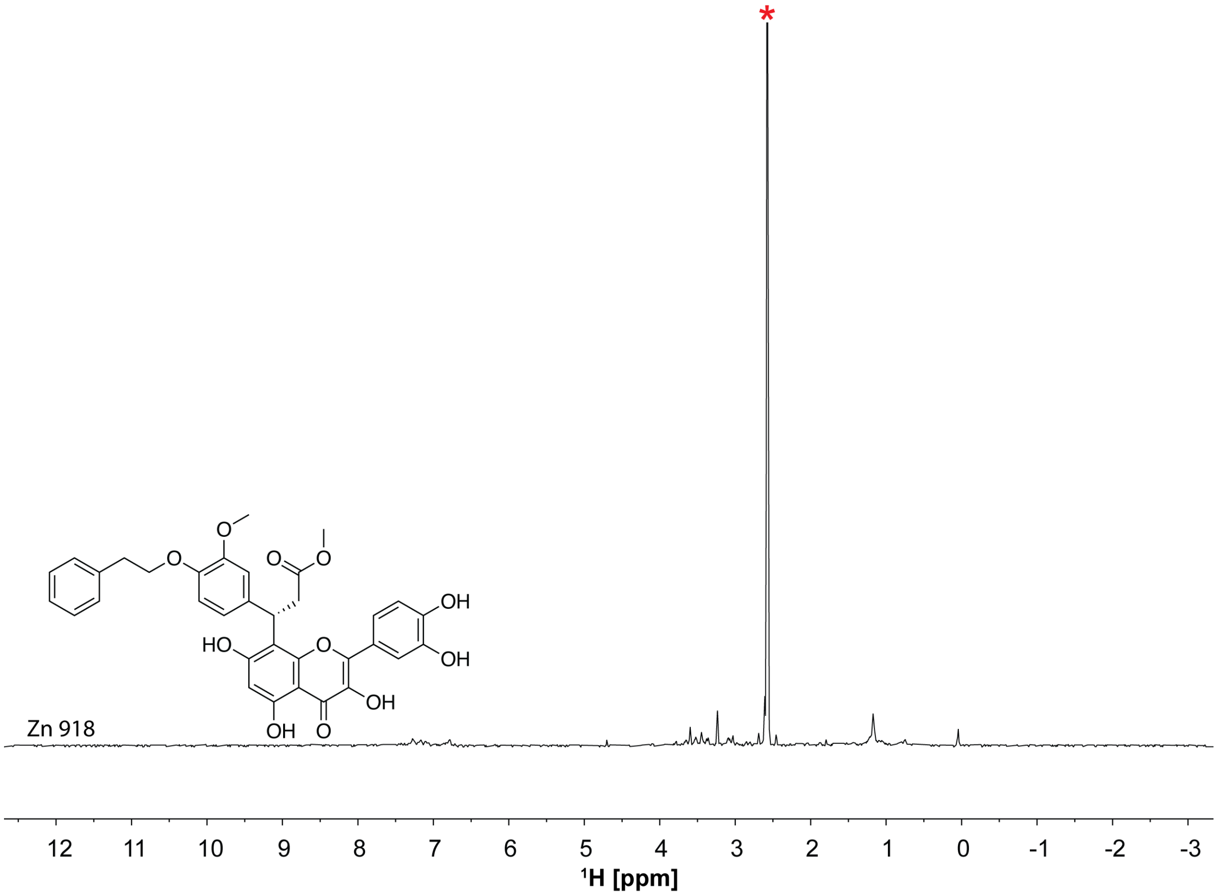


**Figure S21.** Chemical structure and ¹H spectrum of Zn 918. The NMR spectrum was recorded on Zn 918 solubilized in buffer containing 50 mM $KH_2PO_4$, pH = 7.5, 50 mM KCl, 1 mM $MgCl_2$, 95% $D_2O$ and 5% DMSO on a 600 MHz Bruker spectrometer equipped with a cryoprobe. The red asterisk indicates the DMSO peak.

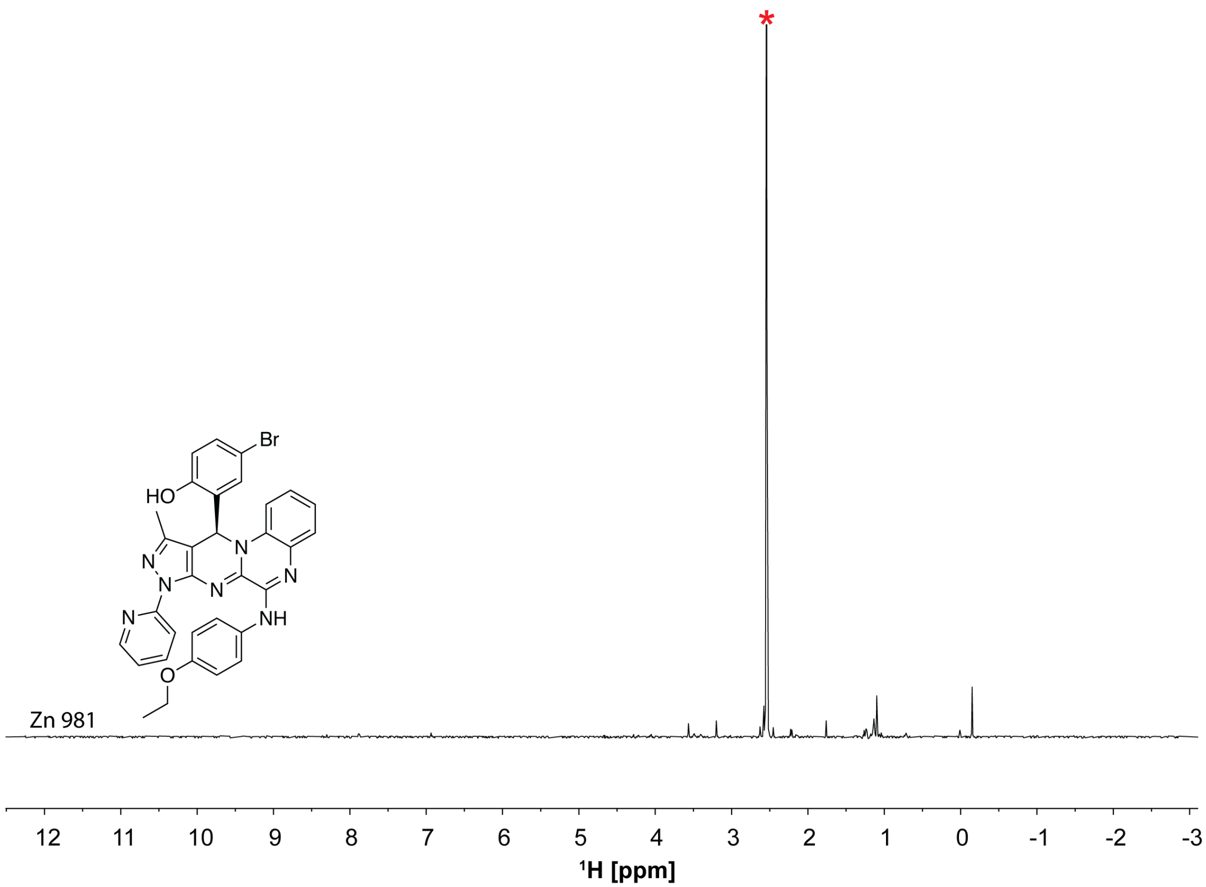


**Figure S22.** Chemical structure and $^1$H spectra of Zn 981. The NMR spectrum was recorded on Zn 981 solubilized in buffer containing 50 mM $KH_2PO_4$, pH = 7.5, 50 mM KCl, 1 mM $MgCl_2$, 95% $D_2O$ and 5% DMSO on a 600 MHz Bruker spectrometer equipped with a cryoprobe. The red asterisk indicates the DMSO peak.

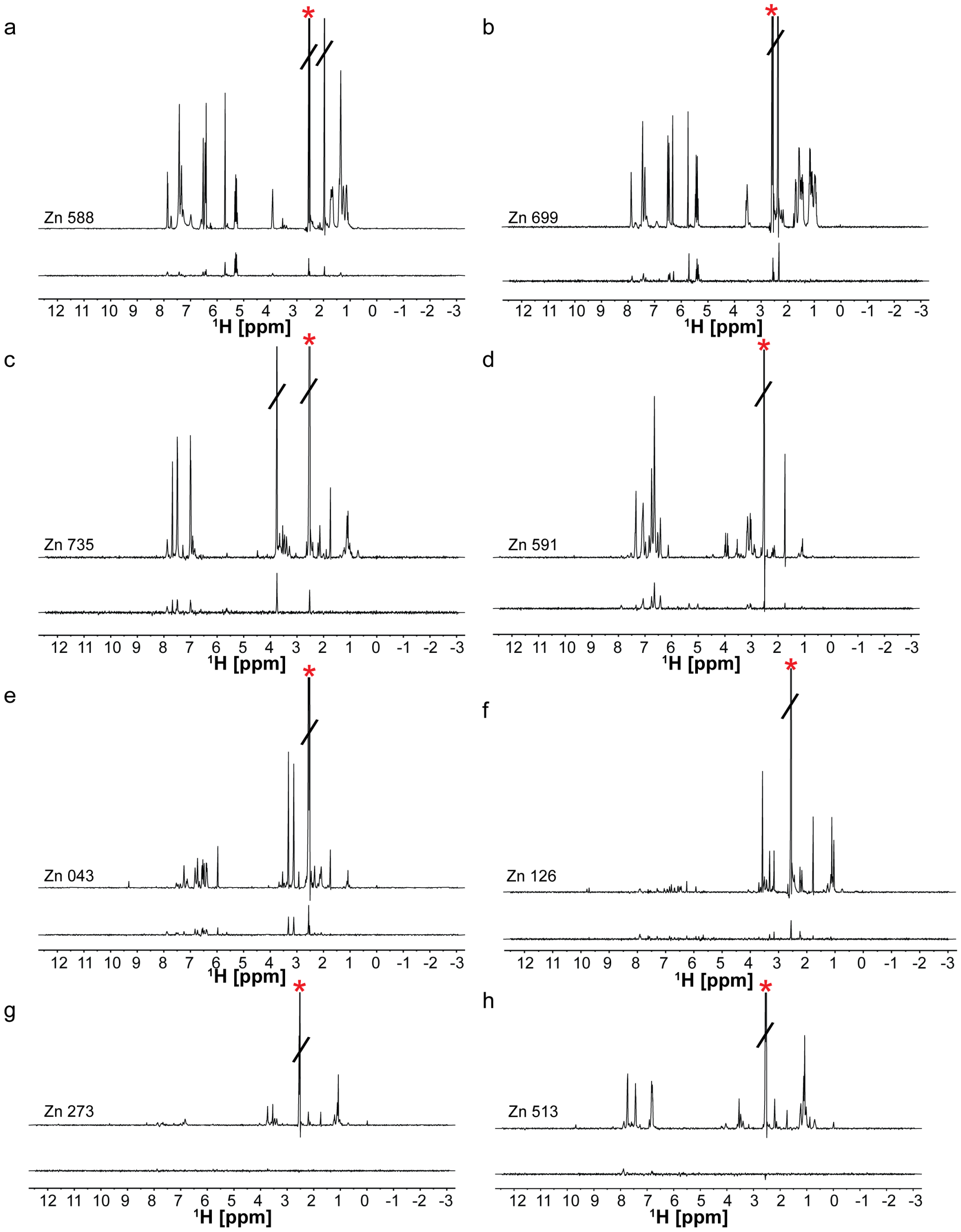


**Figure S23.** STD-NMR based screening and hit identification. The 'off' resonance spectrum (top) and difference spectrum (bottom) are shown for (a) Zn 588 (b) Zn 699 (c) Zn 735 (d) Zn 591 (e) Zn 043 (f) Zn 126 (g) Zn 273 (h) Zn 513. The red asterisk indicates the DMSO peak.

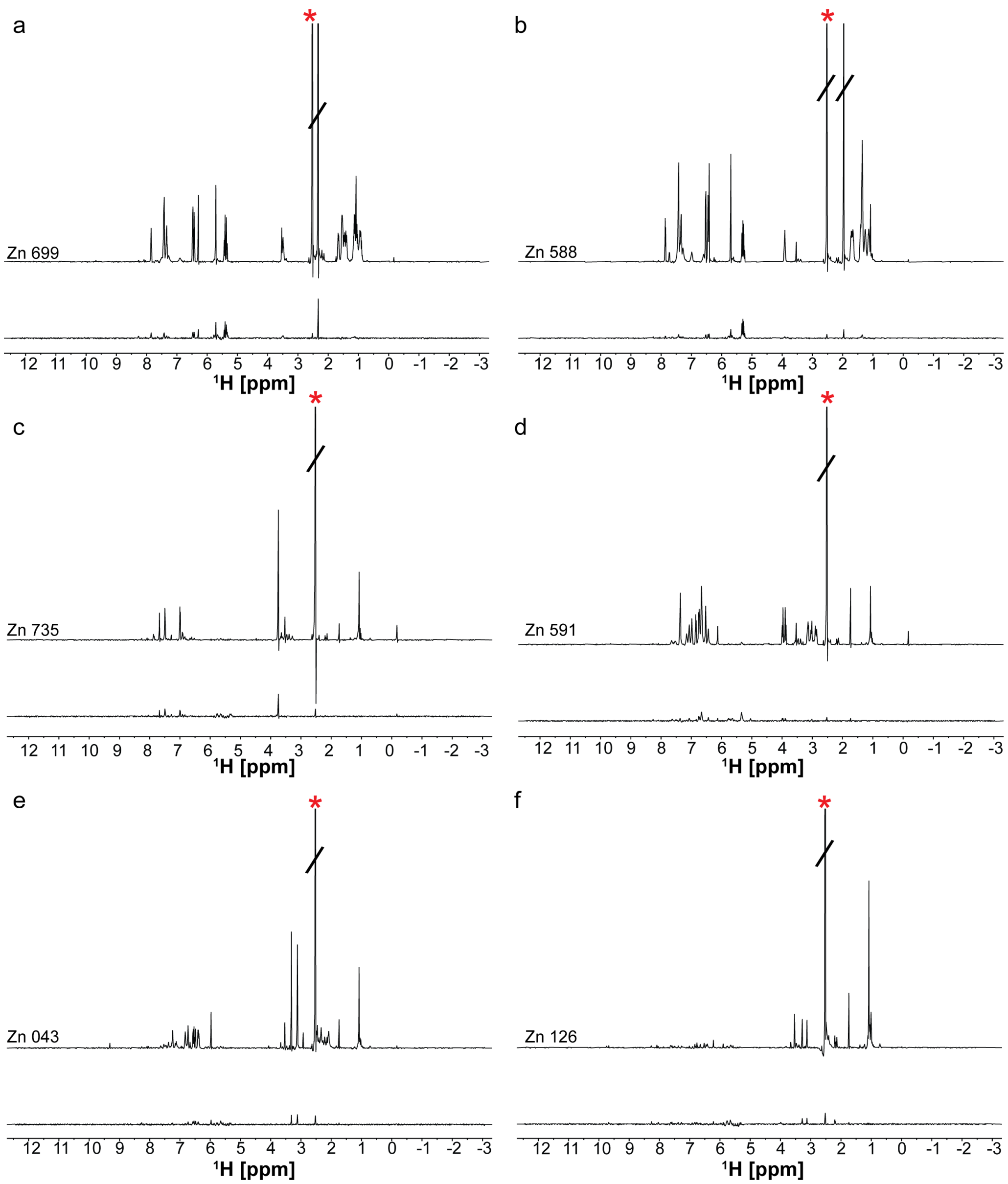


**Figure S24.** STD-NMR screening of NPSL2 hits against TAR RNA. The 'off' resonance spectrum (top) and difference spectrum (bottom) are shown for (a) Zn 699 (b) Zn 588 (c) Zn 735 (d) Zn 591 (e) Zn 043 (f) Zn 126. The red asterisk indicates the DMSO peak.

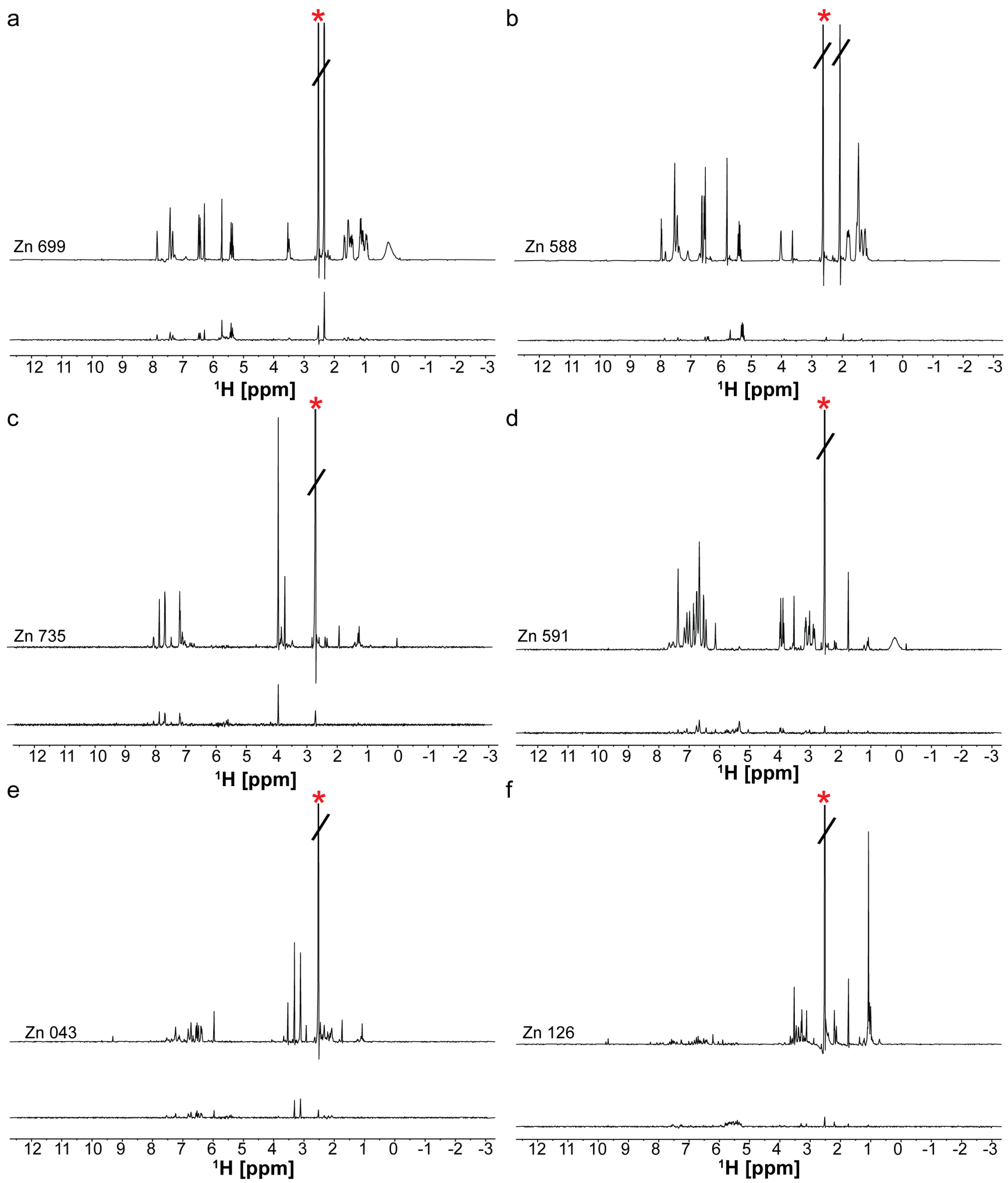


**Figure S25.** STD-NMR screening of NPSL2 hits against Telomerase P2a RNA. The 'off' resonance spectrum (top) and difference spectrum (bottom) are shown for (a) Zn 699 (b) Zn 588 (c) Zn 735 (d) Zn 591 (e) Zn 043 (f) Zn 126. The red asterisk indicates the DMSO peak.

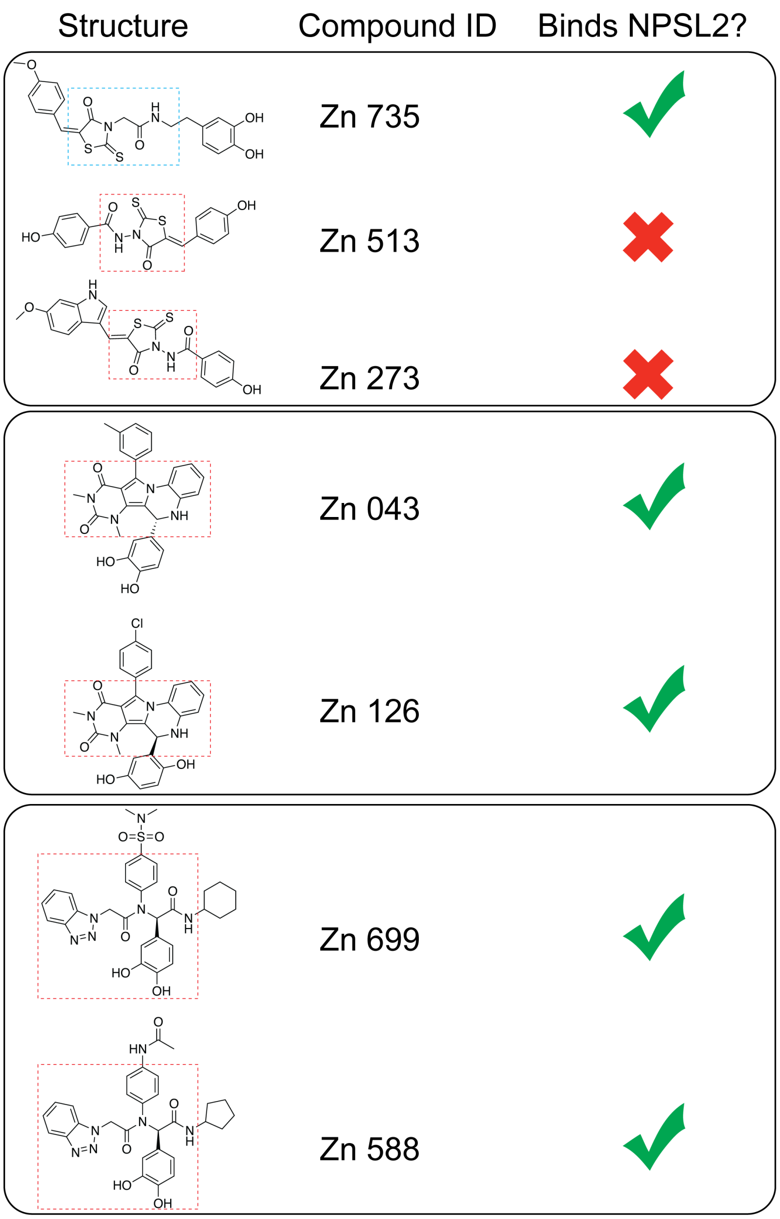


**Figure S26.** Comparison of binding interactions in closely related structures

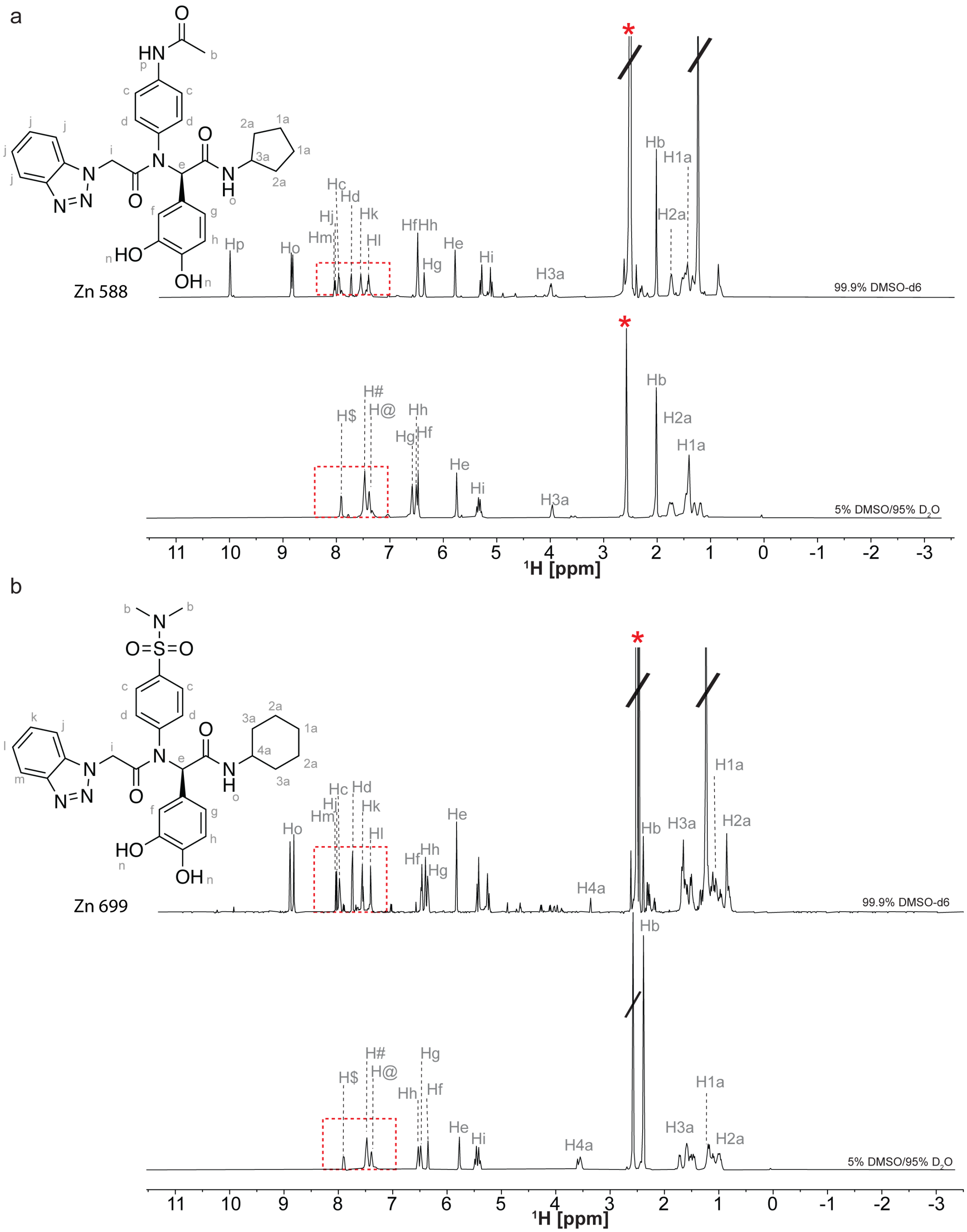


**Figure S27.** Chemical Structure and $^1$H-NMR spectrum of (a) Zn 588 in 99.9% DMSO (top) and 5% DMSO/95% $D_2O$ (bottom). (b) Zn 699 in 99.9% DMSO (top) and 5% DMSO/95% $D_2O$ (bottom). Chemical shifts were assigned using CHEMDRAW. The red asterisk indicates the DMSO peak.

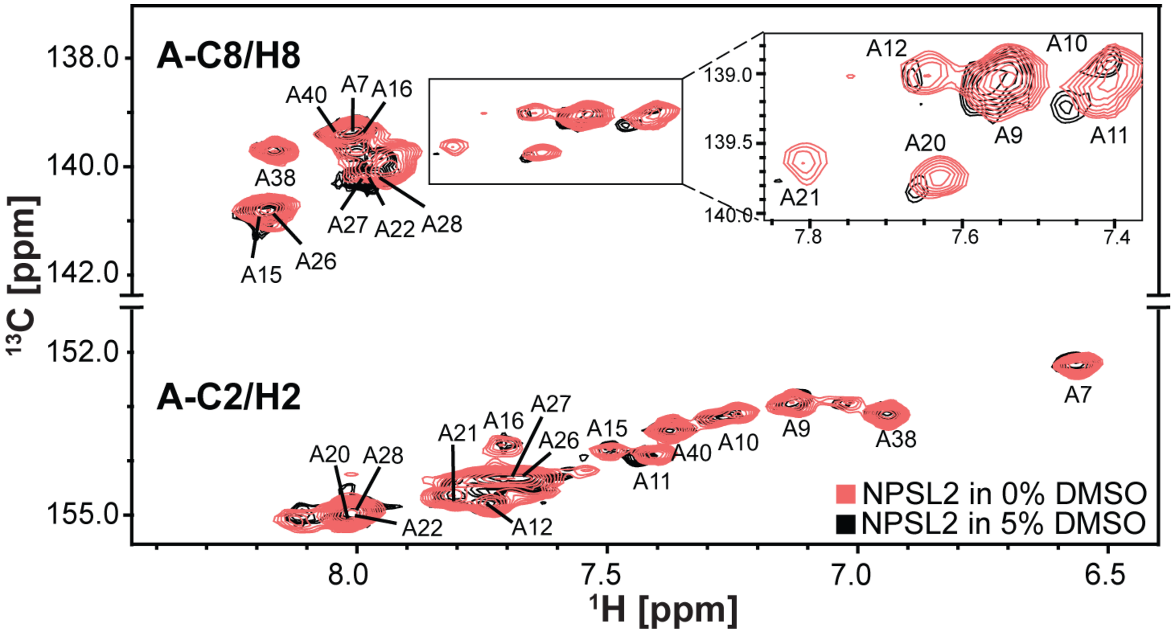


**Figure S28.** HSQC spectra of NPSL2 at 0% DMSO (v/v) (red) and 5% (v/v) DMSO (black).

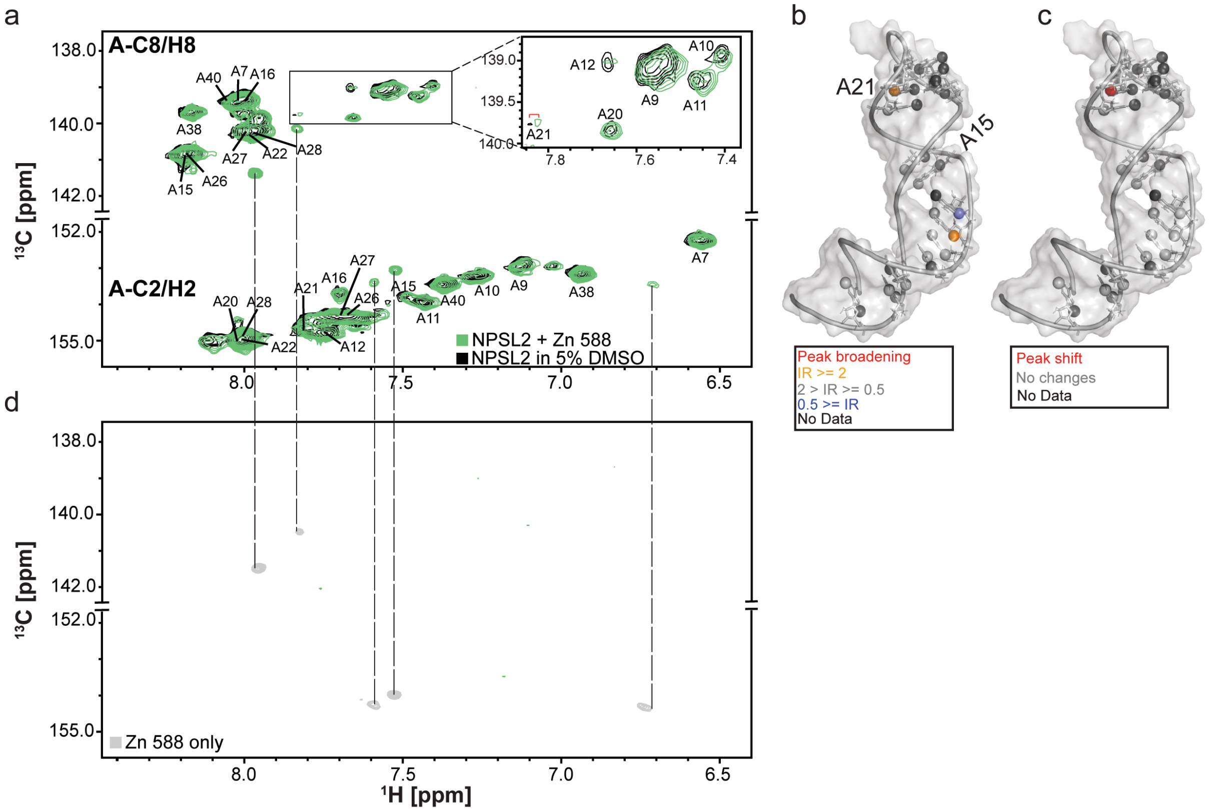


**Figure S29.** NMR chemical shift mapping experiments. (a) HSQC spectrum of NPSL2 alone (black) and with Zn 588 (green). (b) Changes in peak intensities (C2-H2 and C8-H8) that result from ligand binding are mapped onto the NPSL2 structure. (c) Changes in peak positions upon ligand binding are mapped onto the NPSL2 structure. All annotations highlighted in black represent no quantification due to signal overlap. (d) HSQC spectrum of Zn 588. All residues showing shifts in peak positions are indicated with a red bracket and residues showing perturbations arising from complete signal broadening are marked with a red circle on the spectra

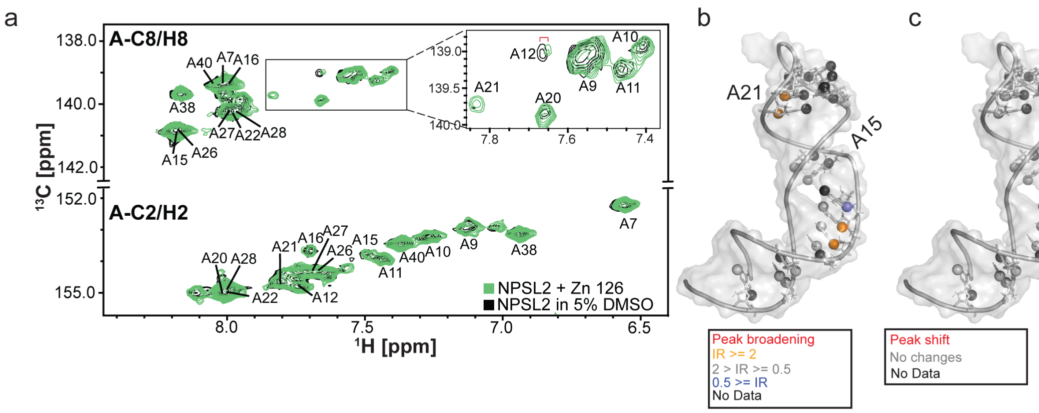


**Figure S30.** NMR chemical shift mapping experiments. (a) HSQC spectrum of NPSL2 alone (black) and with Zn 126 (green). (b) Changes in peak intensities (C2-H2 and C8-H8) that result from ligand binding are mapped onto the NPSL2 structure. (c) Changes in peak positions upon ligand binding are mapped onto the NPSL2 structure. All annotations highlighted in black represent no quantification due to signal overlap. All residues showing shifts in peak positions are indicated with a red bracket and residues showing perturbations arising from complete signal broadening are marked with a red circle on the spectra

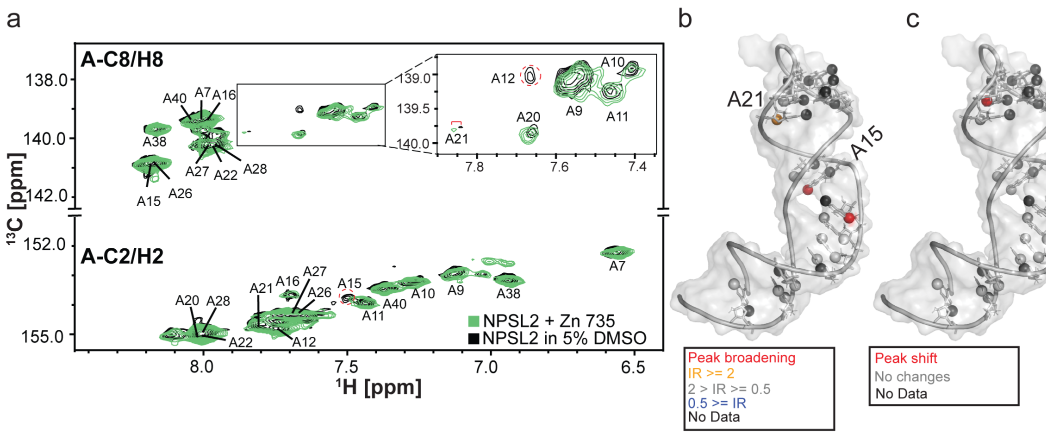


**Figure S31.** NMR chemical shift mapping experiments. (a) HSQC spectrum of NPSL2 alone (black) and with Zn 735 (green). (b) Changes in peak intensities (C2-H2 and C8-H8) that result from ligand binding are mapped onto the NPSL2 structure. (c) Changes in peak positions upon ligand binding are mapped onto the NPSL2 structure. All annotations highlighted in black represent no quantification due to signal overlap. All residues showing shifts in peak positions are indicated with a red bracket and residues showing perturbations arising from complete signal broadening are marked with a red circle on the spectra

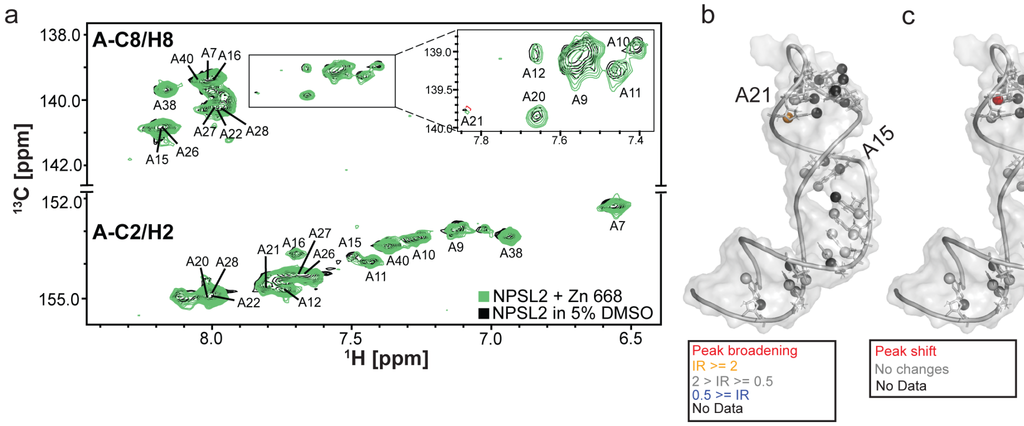


**Figure S32.** NMR chemical shift mapping experiments. (a) HSQC spectrum of NPSL2 alone (black) and with Zn 668 (green). (b) Changes in peak intensities (C2-H2 and C8-H8) that result from ligand binding are mapped onto the NPSL2 structure. (c) Changes in peak positions upon ligand binding are mapped onto the NPSL2 structure. All annotations highlighted in black represent no quantification due to signal overlap. All residues showing shifts in peak positions are indicated with a red bracket and residues showing perturbations arising from complete signal broadening are marked with a red circle on the spectra

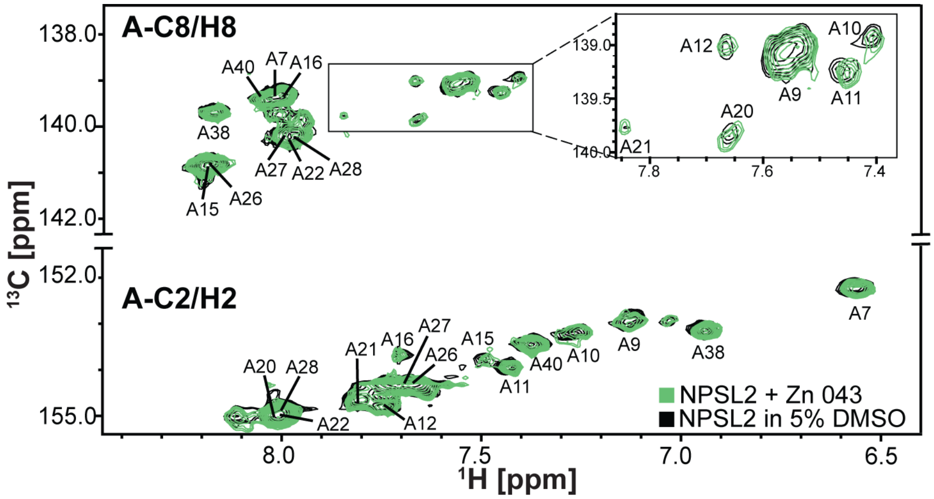


**Figure S33.** Overlayed HSQC spectra of NPSL2 alone (black) and with Zn 043 (green).

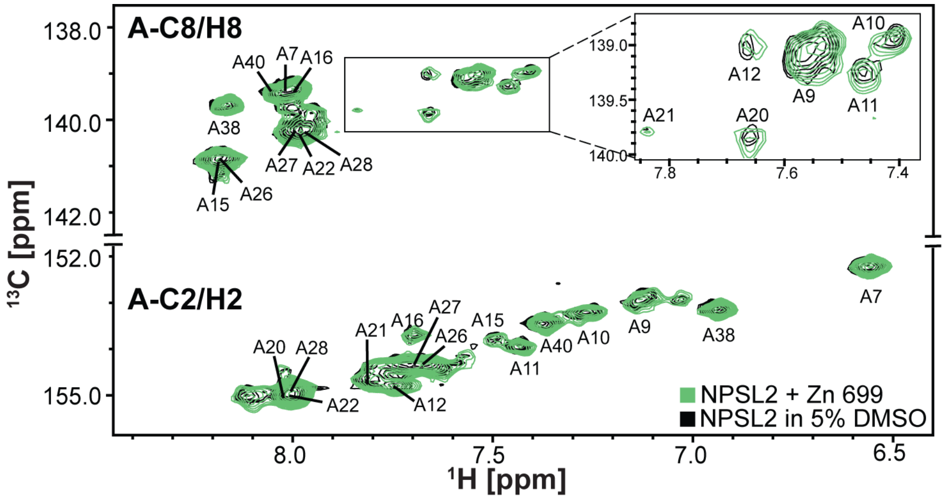


**Figure S34.** Overlayed HSQC spectra of NPSL2 alone (black) and with Zn 699 (green).

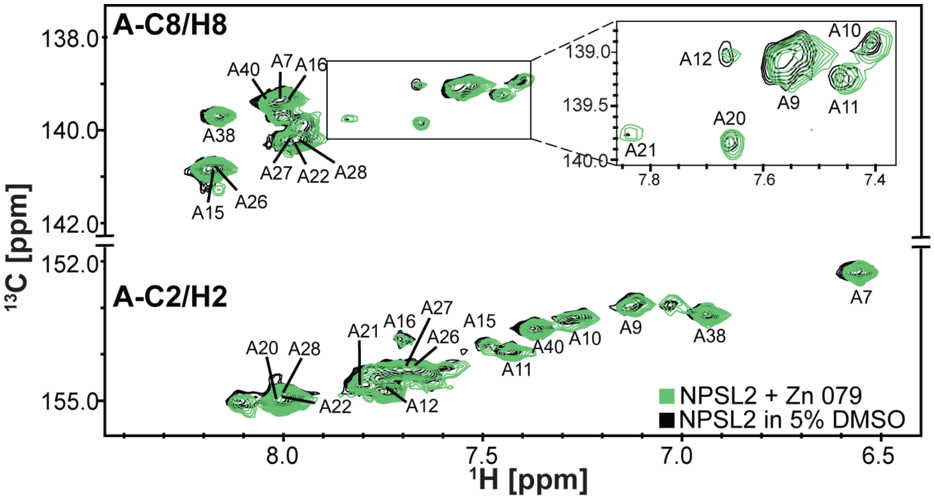


**Figure S35.** Overlayed HSQC spectra of NPSL2 alone (black) and with Zn 079 (green).

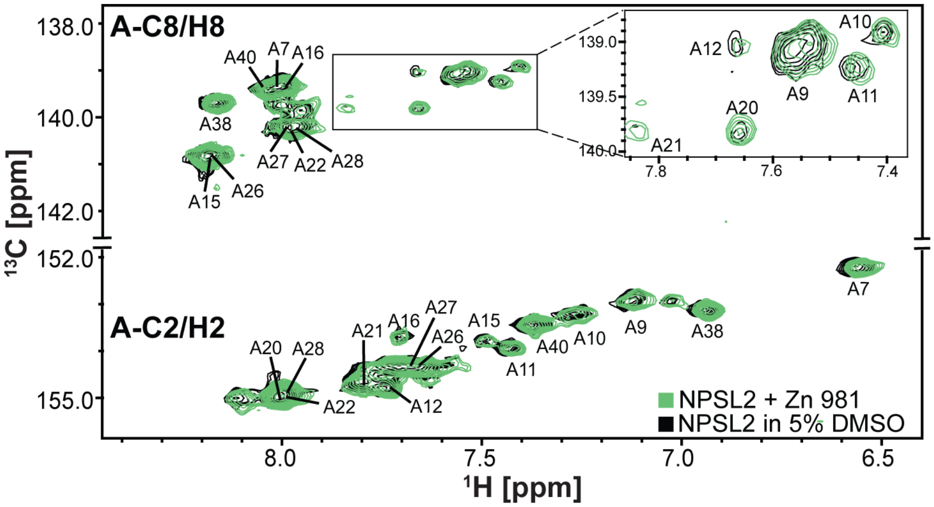


**Figure S36.** Overlayed HSQC spectra of NPSL2 alone (black) and with Zn 981 (green).

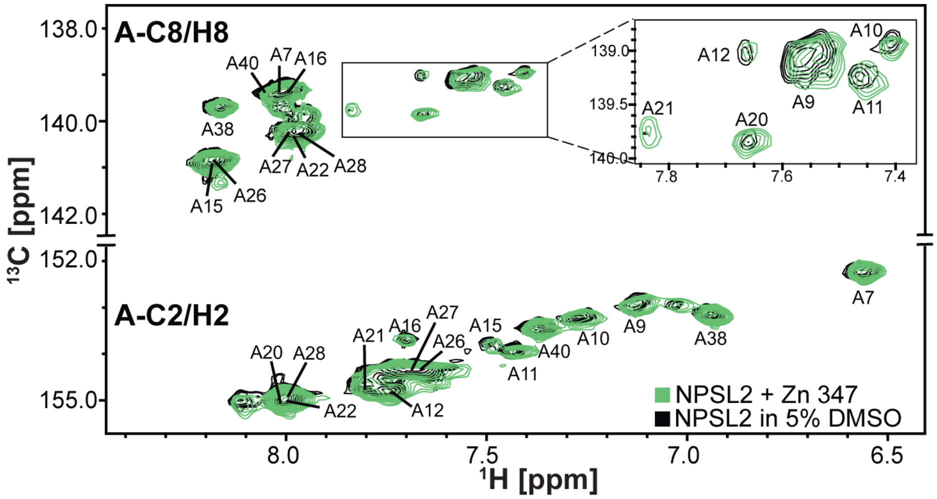


**Figure S37.** Overlayed HSQC spectra of NPSL2 alone (black) and with Zn 347 (green).

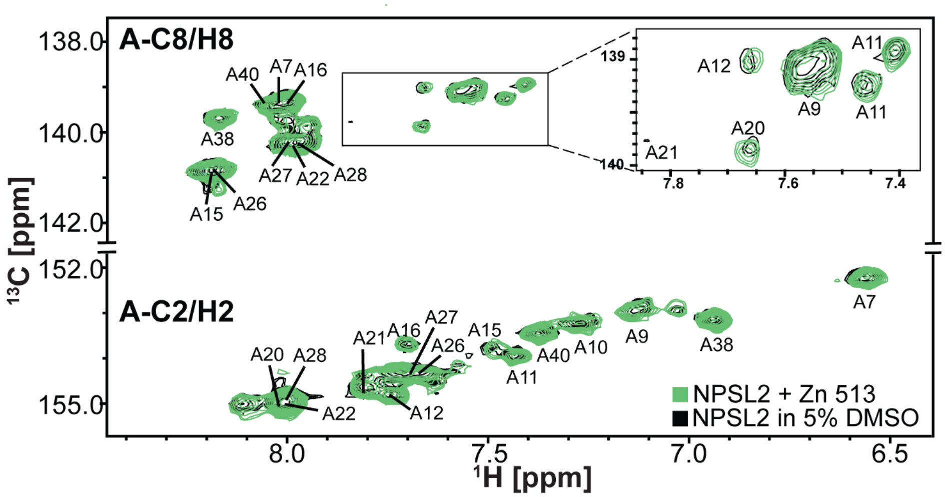


**Figure S38.** Overlayed HSQC spectra of NPSL2 alone (black) and with Zn 513 (green).

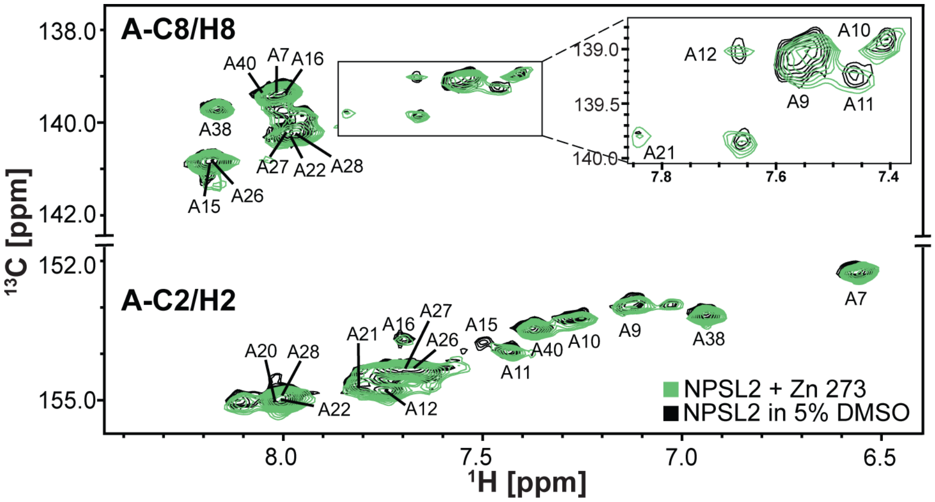


**Figure S39.** Overlayed HSQC spectra of NPSL2 alone (black) and with Zn 273 (green).

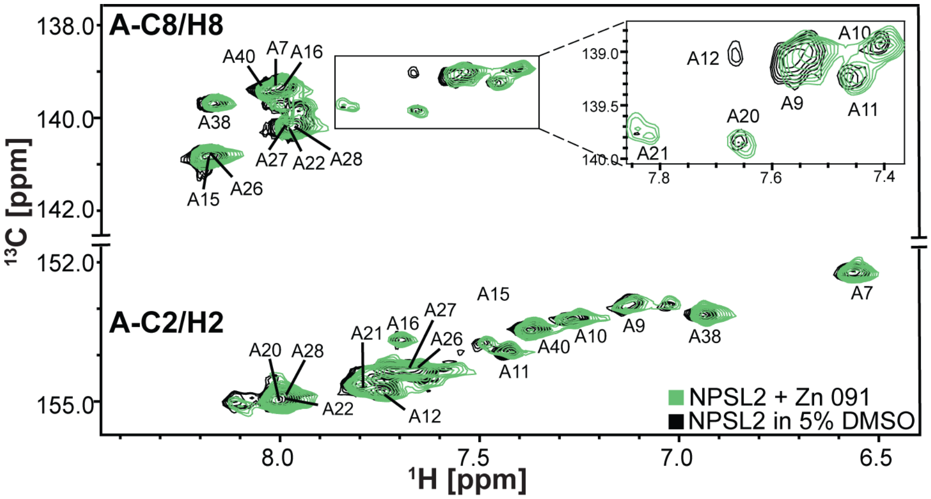


**Figure S40.** Overlayed HSQC spectra of NPSL2 alone (black) and with Zn 091 (green).

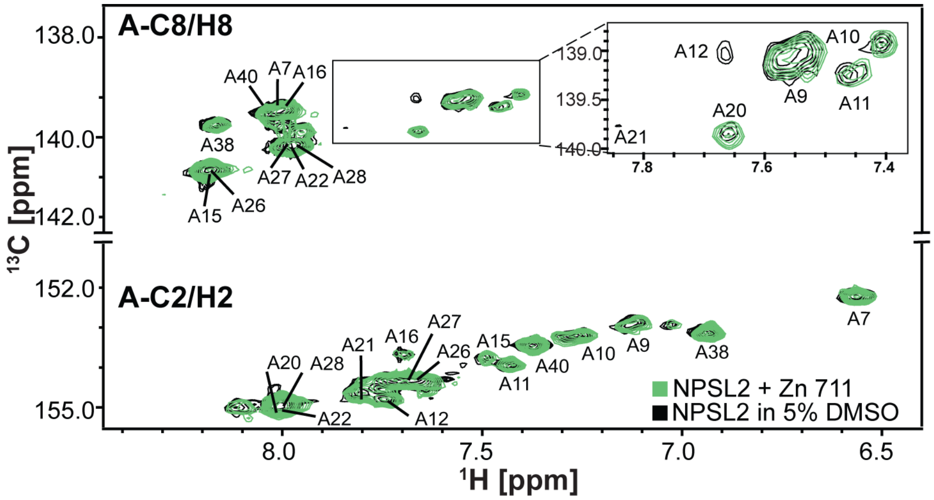


**Figure S41.** Overlayed HSQC spectra of NPSL2 alone (black) and with Zn 711 (green).

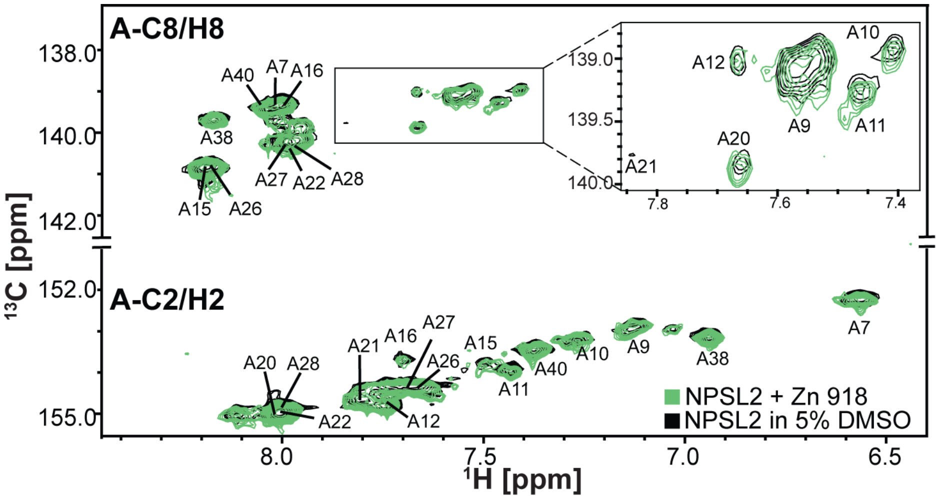


**Figure S42.** Overlayed HSQC spectra of NPSL2 alone (black) and with Zn 918 (green).

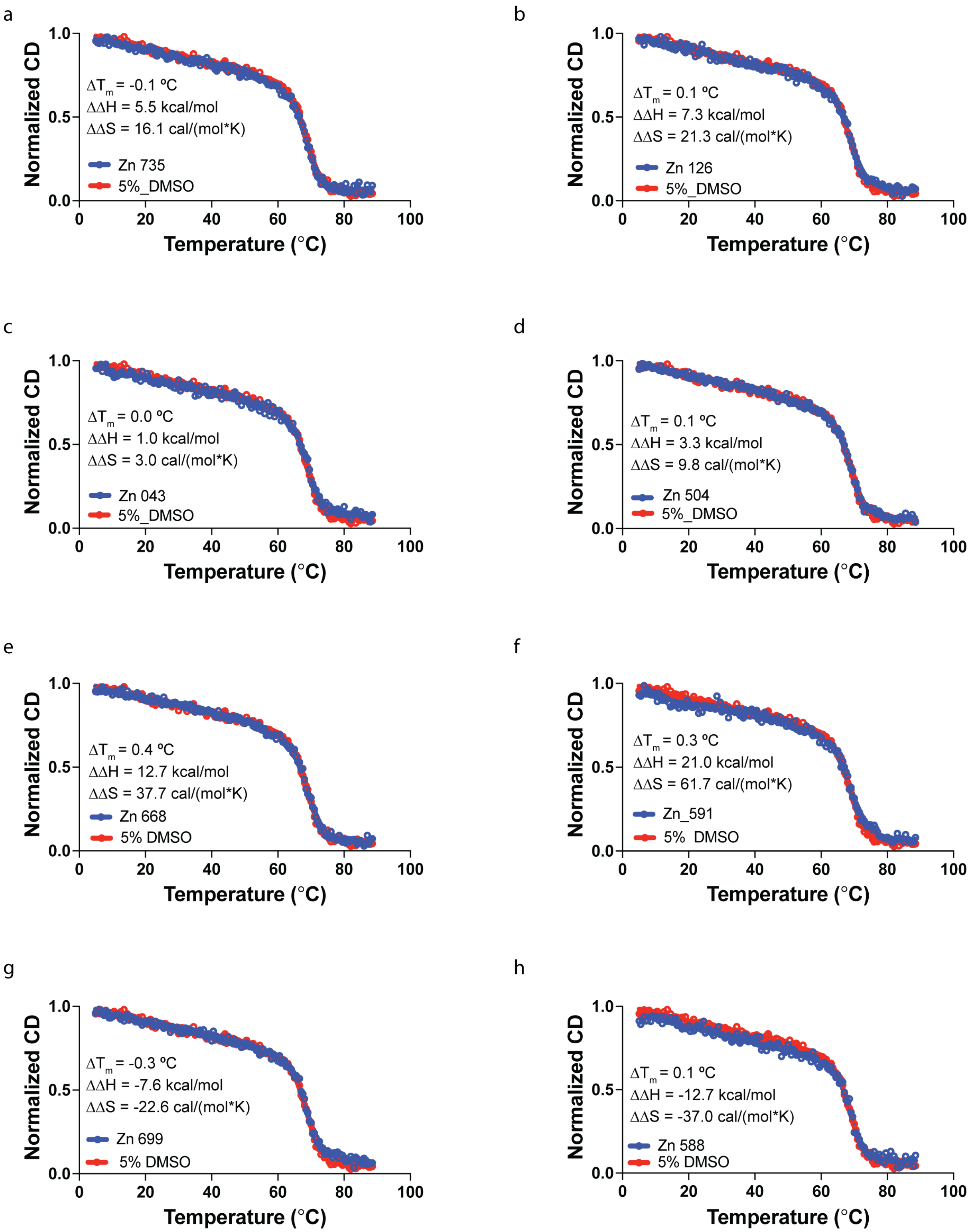


**Figure S43**. Normalized CD thermal denaturation curves for NPSL2 in 5% DMSO (red) and NPSL2 in the presence of ~ 200 μM (a) Zn 735, (b) Zn 126, (c) Zn 043, (d) Zn 504, (e) Zn 668, (f) Zn 591, (g) Zn 699, and (h) Zn 588.
